# Sub-functionalization and epigenetic regulation of a biosynthetic gene cluster in *Solanaceae*

**DOI:** 10.1101/2024.10.02.615186

**Authors:** Santiago Priego-Cubero, Eva Knoch, Zhidan Wang, Saleh Alseekh, Karl-Heinz Braun, Philipp Chapman, Alisdair Fernie, Chang Liu, Claude Becker

**Affiliations:** Faculty of Biology, Ludwig-Maximilians-Universität München, 82152 Martinsried, Germany; Department of Epigenetics, Institute of Biology, University of Hohenheim, 70599 Stuttgart, Germany; Max-Planck-Institute of Molecular Plant Physiology, 14476 Potsdam-Golm, Germany; Center of Plant Systems Biology and Biotechnology, 4000 Plovdiv, Bulgaria

## Abstract

Biosynthetic gene clusters (BGCs) are sets of often heterologous genes that are genetically and functionally linked. Among eukaryotes, BGCs are most common in plants and fungi, where they ensure the co-expression of the different enzymes coordinating the biosynthesis of specialised metabolites. Here, we report on a newly identified withanolide BGC in *Physalis grisea* (ground-cherry), a member of the nightshade family (*Solanaceae*). A combination of transcriptomic, epigenomic and metabolic analyses, revealed that, following a duplication event, this BGC evolved two tissue-specifically expressed sub-clusters, containing eight pairs of paralogs that contribute to related but distinct biochemical processes; this sub- functionalisation is tightly associated with epigenetic features and the local chromatin environment. The two sub-clusters appear strictly isolated from each other at the chromatin structural level, each forming a highly self-interacting chromatin domain with tissue-dependent levels of condensation. This correlates with gene expression exclusively in either above- or below-ground tissue, thus spatially separating the production of different withanolide compounds. By comparative phylogenomic analysis, we show that the withanolide BGC most likely evolved before the diversification of the *Solanaceae* family and underwent lineage- specific diversifications and losses. The tissue-specific sub-functionalisation is common to species of the *Physalideae* tribe but distinct from other, independent duplication events outside of this clade. In sum, our study reports on the first instance of an epigenetically modulated sub-functionalisation within a BGC and sheds light on the biosynthesis of withanolides, a highly diverse group of steroidal triterpenoids important in plant defence and amenable to pharmaceutical applications due to their anti-inflammatory, antibiotic and anti- cancer properties.

## Introduction

Plants produce a plethora of specialised metabolites that typically serve as protectants, repellents, attractants, or signalling molecules in ecological interactions. Some have growth- stimulating or -inhibitory properties. Among the 200,000 to one million different compounds that plants are estimated to produce, triterpenoids are one of the largest and most diversified groups (Cárdenas et al. 2019). Withanolides, in turn, are highly diverse steroidal triterpenoids that are prominent in *Solanaceae* but have also been detected in some species outside the nightshade family (e.g. Yang et al. (2011)). The name-giving plant species, *Withania somnifera* (commonly referred to by its Sanskrit name “ashwagandha”), has been used as a medicinal plant in traditional Indian medicine for several thousand years; modern pharmaceutical research is investigating several withanolides for their anti-inflammatory properties and their potential use in treatments of cancer and autoimmune diseases (Huang et al. 2020; Zhang et al. 2023; Cigler et al. 2024). Besides in *Withania*, withanolide analogues occur in many but not all *Solanaceae*; they are present in different *Physalis* and *Petunia* species but are notably absent in the majority of *Solanum* species such as tomato and potato. Currently, more than 600 compounds belonging to this class are known, differing in the configuration of their side chains and the positioning of double bonds in their structure (Dhar et al. 2015; Xia et al. 2022). This diversity makes withanolides, which act as herbivore repellents and antibiotics in the plant, a promising source for novel pharmaceutical agents.

The biosynthesis of withanolides branches off the phytosterol pathway. The first dedicated step towards withanolides is mediated by a sterol ι1^24^-isomerase (24ISO) that catalyses the isomerization from 24-methylenecholesterol to 24-methyldesmosterol (Knoch et al. 2018). 24ISO is a homolog of DWARF1 (DWF1), with two paralogs, SSR1 and SSR2 (for Sterol Side- chain Reductase), in *Solanum lycopersicum* (tomato) and *S. tuberosum* (potato). SSR1 catalyses the synthesis of campesterol from 24-methylenecholesterol, an intermediate in brassinolide biosynthesis, while SSR2 plays a role in the production of cholesterol and hence steroidal glycoalkaloids (Sawai et al. 2014; Akiyama et al. 2023).

While 24ISO is the starting point of the withanolide biosynthesis pathway, several enzymatic steps, particularly those typically involving cytochrome P450s (CYP450s) and 2-oxoglutarate- dependent dioxygenases (2ODDs), have been hypothesised as being necessary to complete the biosynthesis from 24-methyldesmosterol to the withanolides (Knoch et al. 2018). Potential gene clusters containing genes encoding CYP450s and 2ODDs and located in the same genomic region as *24ISO* were identified in related solanaceous plants, suggesting a larger withanolide biosynthetic gene cluster (BGC) (Knoch et al. 2018). Several BGCs of plant specialised metabolites have been identified in recent years, e.g. encoding the biosynthesis of diterpenes in rice and barley, or benzoxazinoids in maize and other grasses (Polturak and Osbourn 2021; Polturak et al. 2022). A BGC encoding withanolide production, although hypothesised in the past, has not been identified to date.

Here, we present the discovery of a withanolide BGC in the *Solanaceae* species *Physalis grisea*, a close relative of *W. somnifera*. Our data show that the genes in the cluster play an essential role in the synthesis of withanolides. The cluster architecture revealed a tandem duplication, resulting in a sub-functionalization into two distinct sub-clusters, each containing a set of core genes required for withanolide biosynthesis. Through the combined analysis of transcriptomic, epigenomic and metabolic datasets, we show that sub-cluster expression and withanolide formation are tissue-specific, and that sub-cluster differentiation is associated with tissue-specific epigenetic configuration and higher-order chromatin organisation. Through phylogenomic analysis, we show that this sub-functionalisation is not specific to *P. grisea* but is common to the *Physalideae* tribe. In sum, our study uncovers how biosynthesis of plant specialised metabolites can diversify via duplication and differential organisation of the chromatin environment.

## Results

### The putative withanolide biosynthetic gene cluster in P. grisea underwent tandem duplication

At the time Knoch *et al*. identified 24ISO as the enzyme catalysing the first dedicated biosynthesis step in withanolide production, only a few *Solanaceae* genomes had been assembled and annotated. Nonetheless, the authors had already noticed that the gene encoding the 24ISO ortholog in *Capsicum annuum*, *Solanum melongena* (in which it appeared pseudogenic) and *Petunia inflata* was flanked by genes encoding CYP450s (CYP88 subfamily, among others) and 2-oxoglutarate dependent (2-ODDs) dioxygenases that might be involved in the metabolization of the withanolide-intermediate 24-methyldesmosterol (Knoch et al. 2018). We set out to study the possibility that withanolide biosynthesis was organised in a BGC. In absence of a *W. somnifera* genome assembly, we referred to the most closely related available assembly and gene annotation, that of *P. grisea* (He et al. 2023; Huang et al. 2023). Like *W. somnifera*, *Physalis* spp. belong to the *Physalideae* tribe. Its members produce withanolide analogs, known as physalins (Huang et al. 2020). Given this close phylogenetic and biochemical relationship between the genera *Physalis* and *Withania*, we hypothesised that the *P. grisea* genome could harbour orthologs of *Ws24ISO* and could serve as a proxy to study putative BGCs, particularly since *P. grisea* is currently being established as a new genetic model in the *Solanaceae* family (Van Eck 2022; Dale et al. 2024).

Using the *W. somnifera* 24ISO amino acid sequence as a query for BLAST, we identified two *24ISO* copies in the *P. grisea* genome, localised at the end of chromosome 1 and separated by ∼350 kb (**Figure 1A**). Closer inspection of this genomic region revealed aggregation of several genes encoding enzymes putatively involved in the biosynthesis of secondary metabolites (**Figure 1A**). In line with the observations by Knoch et al. (2018) in *C. annuum*, *S. melongena* and *P. inflata*, we found nine genes that encoded CYP450s belonging to different families and subfamilies (*CYP87G1*, *CYP749B2*, *CYP88C8*, *CYP88C7* and *CYP88C10*), and six *2-ODD* genes surrounding the *24ISO* orthologs, suggesting the existence of a putative withanolide BGC in *P. grisea*. Additionally, we also identified two genes (one of them appearing pseudogenic) encoding Short-Chain Dehydrogenase/Reductases (SDRs), two for *S*ulfotransferases (Sulf), and one encoding an acylsugar acyltransferase (ASAT) (**Figure 1A**). We noticed that eight genes appeared to be duplicated within a 376 kb region, such that overall, the locus exhibited signals of a tandem duplication that had resulted in two groups of genes. Remarkably, each group encoded core paralogs of a putative withanolide biosynthetic pathway (24ISOa and b, respectively, CYP87G1a/b, CYP749B2a/b, CYP88C7a/b, CYP88C10a/b, 2ODD3a/b, Sulfa/b, and SDRa/b), and was separated from the other group by an intermediate gene-depleted region of approximately 50 kb in length (**Figure 1A**). Sequence analysis of the respective gene and protein copies corroborated the notion that these gene pairs were paralogs that most likely had arisen from a duplication, with pairwise nucleotide and amino acid sequence identities ranging between 84.4-94.6% and 80-95%, respectively (**Supplementary Table 1**).

**Figure 1.**
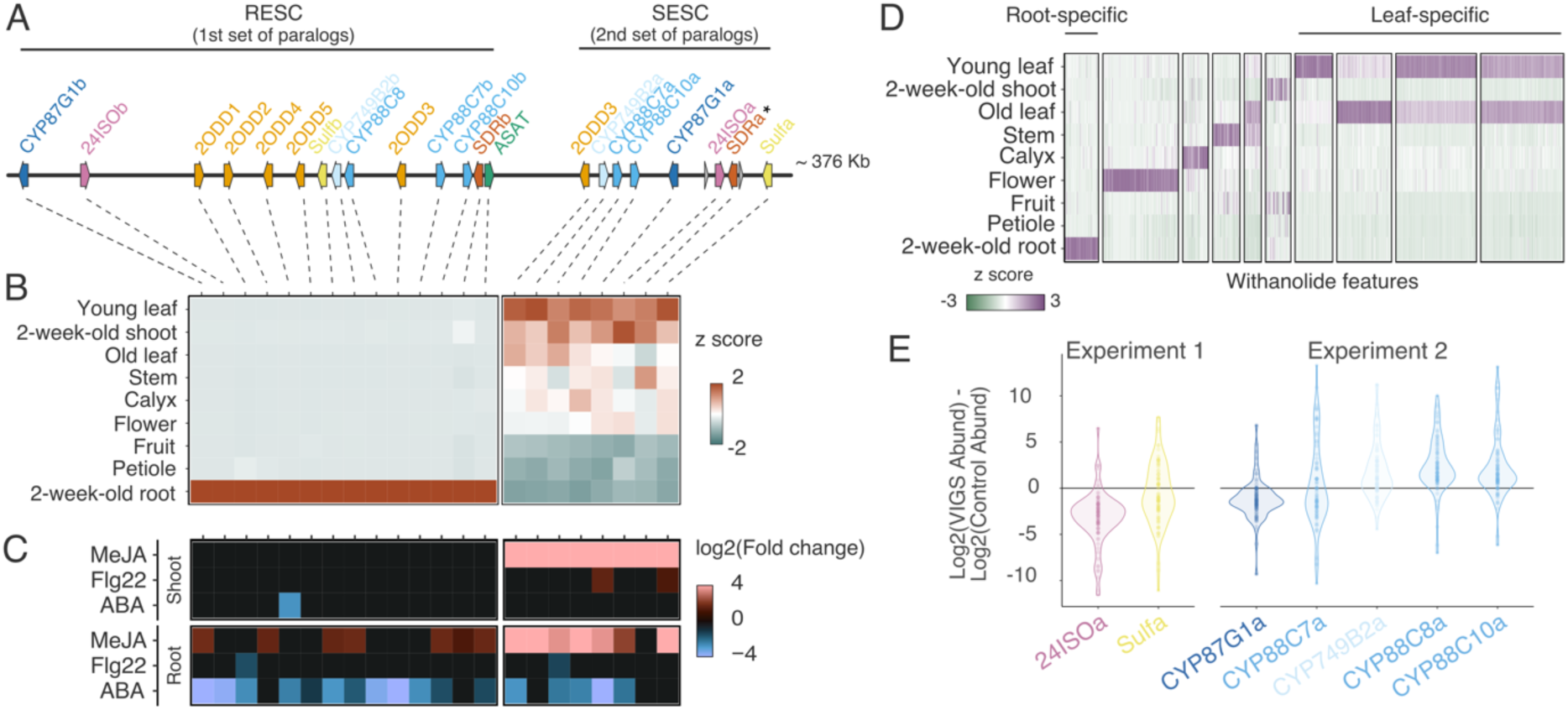
The withanolide biosynthetic gene cluster is organised into two sub-clusters. **A)** Genomic organisation of the withanolide BGC on *P. grisea* chromosome 1; distances between genes are drawn to scale; arrows indicate the sense of transcription. RESC and SESC: root- and shoot-expressed sub-clusters, respectively. Asterisk indicates a paralog that appears to be pseudogenic. **B)** Heatmap showing the relative expression level of each gene within the cluster across different tissues. Expression levels are presented as z-scores of transcripts per million (TPM), relative to the mean for each gene. **C)** Heatmap showing the expression of each gene within the cluster in response to the putative elicitors methyl jasmonate (MeJA), flagellin 22 (Flg22), and abscisic acid (ABA). Differentially expressed genes were defined as those with log2(fold-change) >1 or <(-1) and an adjusted p-value <0.01. Log2(fold-change) values between -1 and 1 were set to black in the colour key. Differential expression was calculated independently for each tissue using the respective mock treatment as the reference. **D)** Relative abundance of withanolides across the same tissues as in (B), expressed as the z-score of the intensity values relative to the mean for each compound. **E**) Relative abundance of the leaf-specific withanolides in (D) after VIGS of different genes from the shoot-expressed sub-cluster.

In sum, our data showed that *P. grisea* has two *24ISO* paralogs that are embedded in a putative withanolide BGC and which appeared to have undergone a partial tandem duplication. This raised the questions whether this cluster architecture was specific to *P. grisea* or widespread in the *Solanaceae* family, and whether the tandem duplication had any functional consequences.

### Withanolide production is organised by two differentially regulated sub-clusters

A key feature of BGCs is the co-regulation of expression of the genes that form the cluster, which is thought to ensure the faithful spatio-temporal production of a given metabolite, also in response to different elicitors (Nützmann and Osbourn 2015). To test whether the members of the putative withanolide BGC were co-regulated, we performed whole-genome transcriptomic analysis via mRNA sequencing (RNA-seq) across two sets of samples (**Supplementary Dataset 1**). The first consisted of nine different tissues and developmental stages: shoot and root tissue of two-week-old seedlings; young leaf, mature leaf, flower, stem, petiole, calyx, and fruit (**Supplementary Figure s1**). For the second set, we extracted RNA from roots and shoots of two-week-old seedlings treated with methyl jasmonate (MeJA), which is known to trigger withanolide production in *W. somnifera* (Rana et al. 2014). We further included treatments with the stress hormone abscisic acid (ABA), which induces expression of the withanolide analogue transport protein PDR2 in *Petunia* (Sasse et al. 2016), and the bacterial elicitor flagellin 22 (Flg22) to explore their respective impact on the gene cluster expression.

We combined the data from both RNA-seq experiments in a co-expression analysis using weighted gene correlation network analysis (WGCNA) (Zhang and Horvath 2005; Langfelder and Horvath 2008; Russo et al. 2018) (**Supplementary Figure 2**). Instead of the expected co-regulation of all the genes in the putative BGC, the two groups of paralogs showed distinct and contrasting tissue-specific expression: the first 14 collinear genes, containing the first set of core paralogs in the BGC (**Figure 1A**), were exclusively expressed in root (expression module M2) (**Supplementary Figure 2D**). In contrast, the second set of eight paralogs were all placed in module M1, which grouped genes with high expression in shoot and aerial tissues (**Figure 1B; Supplementary Figure 2D**). Based on these expression profiles, we named these sub-clusters ‘root-expressed sub-cluster’ (RESC) and ‘shoot-expressed sub-cluster’ (SESC), respectively.

In the treatment data, the expression of each sub-cluster was elicited by MeJA. Genes from the SESC responded in both shoot and root tissue, while the RESC genes responded exclusively in root tissue, indicating that the expression of the RESC is tightly repressed in shoot tissue. In contrast, ABA repressed the expression of both the RESC and the SESC in roots (**Figure 1C; Supplementary Figure 2F**); neither sub-cluster responded to flg22. Our data indicate that the tandem duplication resulted in contrasting expression patterns of the two sub-clusters, which seem to be regulated tissue-specifically not only in their respective constitutive expression but also in their response to external stimuli.

Duplication of enzymes in specialised metabolism often leads to neofunctionalization and, in turn, the production of novel metabolites (Smit and Lichman 2022). We hypothesised that the sub-division and tissue-specific regulation of expression could indicate the sub- functionalization of the putative withanolide BGC and result in qualitative differences in withanolide production in the respective tissues. To test this hypothesis, we performed untargeted metabolomics on the tissue samples used for the RNA-seq. After several filtering steps to select features potentially derived from withanolides in the MS/MS spectrum, we compiled a list of withanolide-associated features (see *Material and Methods* for details). We observed a distinct pattern of tissue-specific accumulation of different withanolide-related features in roots and diverse aerial tissues (**Figure 1D**). To further confirm the role of the BGC in withanolide biosynthesis, we employed virus-induced gene silencing (VIGS). Since VIGS can only be applied in leaves but not in roots, we targeted genes from the SESC. The silencing of *24ISOa*, *CYP87G1a* and *Sulfa* led to an overall decrease in the abundance of leaf-specific withanolide features (**Figure 1E; Supplemental Figure 3**). Silencing of *CYP88C7a* had similar effects, but we also noted higher levels for some features, which potentially reflects an accumulation of certain intermediates (**Figure 1E**). Lastly, silencing of *CYP749B2a*, *CYP88C8a* and *CYP88C10a* led to the reduction of few withanolide features and the general accumulation of other, potentially intermediate, products (**Figure 1E**).

Taken together, these results provide support for the existence of two functionally distinct sub- clusters that produce a distinct array of withanolides in a tissue-specific manner.

### The withanolide sub-clusters are functionally isolated from each other and display distinct levels of tissue-specific chromatin compaction

The presumed tandem duplication resulted in two neighbouring gene clusters with highly distinct expression profiles, raising the question how such a functional division could be organised and maintained at the regulatory level. Given the size of the sub-clusters and the number of genes (14 and 8, respectively) that were coordinately expressed, we hypothesised that canonical transcription-factor-mediated regulation might insufficiently explain the observed pattern. Several recent studies have highlighted that local chromatin accessibility and higher-order organisation at the three-dimensional (3D) level can play an important role in the coordinated regulation of gene expression of BGCs in plants (Nützmann et al. 2020; Yang et al. 2021; Li et al. 2023). BGCs are located within topologically associated domains (TADs), self-interacting chromatin regions with suppressed interaction with neighbouring chromatin (Doğan and Liu 2018), or within regions with TAD-like features (Nützmann et al. 2020). This topological organisation facilitates precise spatio-temporal co-expression of the genes within the cluster, enabling long-range interactions between genes and between genes and their *cis*-regulatory elements (Zhao et al. 2022; Li et al. 2023).

To test whether the distinct expression patterns of the RESC and SESC are regulated at the chromatin level, we first performed Assay for Transposase-Accessible Chromatin using sequencing (ATAC-seq) of *P. grisea* root and leaf tissue, in four biological replicates per tissue (**Supplementary Figures 4-6**; **Supplementary Dataset 2**). Our data clearly showed that local chromatin accessibility in the RESC was high in root tissue, but that the chromatin in the same sub-cluster was inaccessible in leaves (**Figure 2A** and **2B**). We observed the opposite for the SESC: while the ATAC-seq showed many open chromatin regions in leaf tissue, the same region was less accessible in roots. Together, these results corroborated our hypothesis that the chromatin landscape differed between tissues; in general, each sub-cluster was more accessible in the tissue in which it was expressed. Additionally, we observed that most of the peaks in both tissues and sub-clusters were located in intergenic regions, suggesting a role of distal *cis*-regulatory elements and long-range chromatin contacts to the tissue-specific regulation of each sub-cluster (**Figure 2C**).

**Figure 2.**
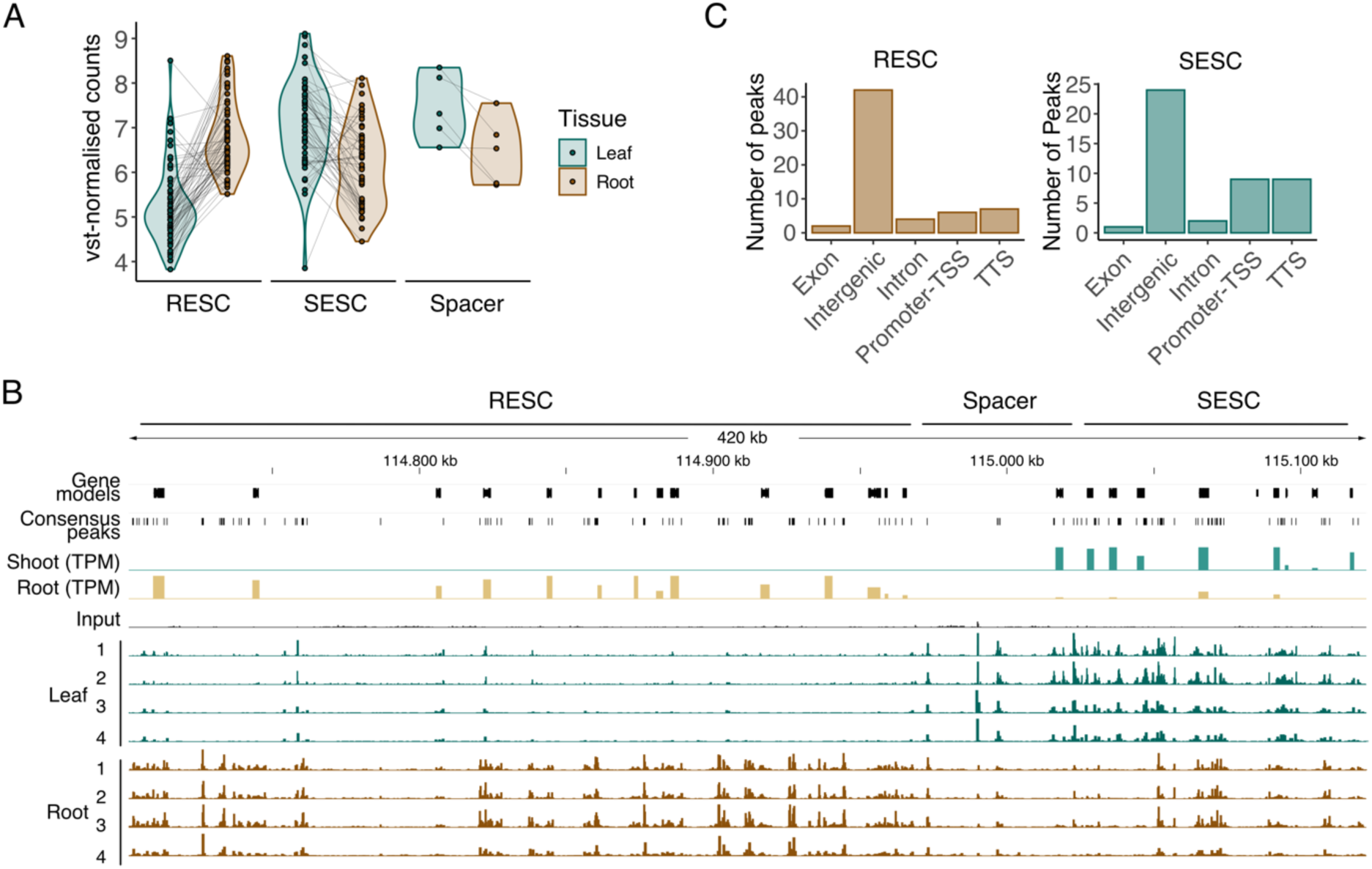
**Tissue-specific local chromatin accessibility at the locus of the withanolide BGC by ATAC-seq**. **A)** Normalised read counts after variance-stabilising transformation using the DESeq2 package (Love et al. 2014) of the consensus narrow peaks called on the Tn5-sensitive regions within each subcluster for a combination of four replicates per tissue. Black lines connect the value of each narrow peak in leaf and root for each subcluster. **B)** Tn5 integration footprint in the withanolide BGC region. From top to bottom: gene models, consensus narrow peaks, expression (TPM: transcript per million; scale was set to 0-400 for both tracks) of 2-week-old shoot and root tissue. ATAC-seq profiles are represented as reads per million (RPM; scale was set to 0-1 for all tracks) for input, leaf (four replicates) and root (four replicates) samples. **C)** Number of narrow peaks in the root and shoot subclusters, separately called for root and leaf samples, respectively, and their location with respect to the genes on each subcluster. Promoter-TSS = promoter and transcription start site, defined from -1 kb to +100 bp; TTS = transcription termination site, defined from -100 bp to +1 kb.

To gain further insights into the chromatin configuration at this genomic locus, we captured whole-genome chromatin-chromatin interactions in *P. grisea* roots and leaves using Hi-C (Lieberman-Aiden et al. 2009) (**Supplementary Dataset 3**). We hypothesised that each subcluster would exhibit distinct patterns of chromatin interactions in a tissue-specific manner. We obtained a total of ∼103 million valid unique read pairs, combining data from two independent biological replicates for each tissue (see *Material and Methods* for details; **Supplementary Figure 7A**). We first examined the genome packaging of *P. grisea* at the chromosomal level with a Hi-C contact map at 1 Mb resolution (**Supplementary Figure 7B**).

The Hi-C map revealed prominent interactions along the main diagonal, attributed to the increased likelihood of interactions among neighbouring linear genome segments (**Supplementary Figure 7B**). We also detected intense intra- and interchromosomal interactions at the chromosomal ends (**Supplementary Figure 7B**), where gene density is highest in the *P. grisea* genome (**Supplementary Figure 7C**). In an A/B compartment annotation of each chromosome, which generally classifies open, euchromatic vs. more compacted, heterochromatic regions, those gene-rich regions at the distal chromosome ends were distinct from the large central region, where transposon density is highest (**Supplementary Figure 7C**) (He et al. 2023). Consequently, we annotated the distal ends as A (accessible) and the central part as B (inaccessible) compartments. This chromosomal organisation was highly similar to that observed in tomato (*S. lycopersicum*) (Dong et al. 2017).

We next explored the withanolide BGC locus, located at the end of chromosome 1 (114.71- 115.12 Mb), by generating normalised Hi-C maps at 20 kb resolution. In both root and shoot tissue, chromatin interactions between the RESC and SESC were rare, indicating that both subclusters were isolated from each other at the chromatin level (**Figure 3A**). Such a spatial separation may contribute to the regulation of tissue-specific expression, allowing each sub- cluster to interact with specific cis-elements. In contrast, the RESC and, to a lesser extent, the SESC exhibited frequent intra-sub-cluster chromatin interactions (**Figure 3A**). We further noticed that the RESC exhibited more frequent interactions with itself in the shoot compared to the root; a weaker, opposite trend applied to the SESC (**Figure 3A)**. This became more obvious when considering the differential chromatin interactions between tissues (**Figure 3B**).

**Figure 3.**
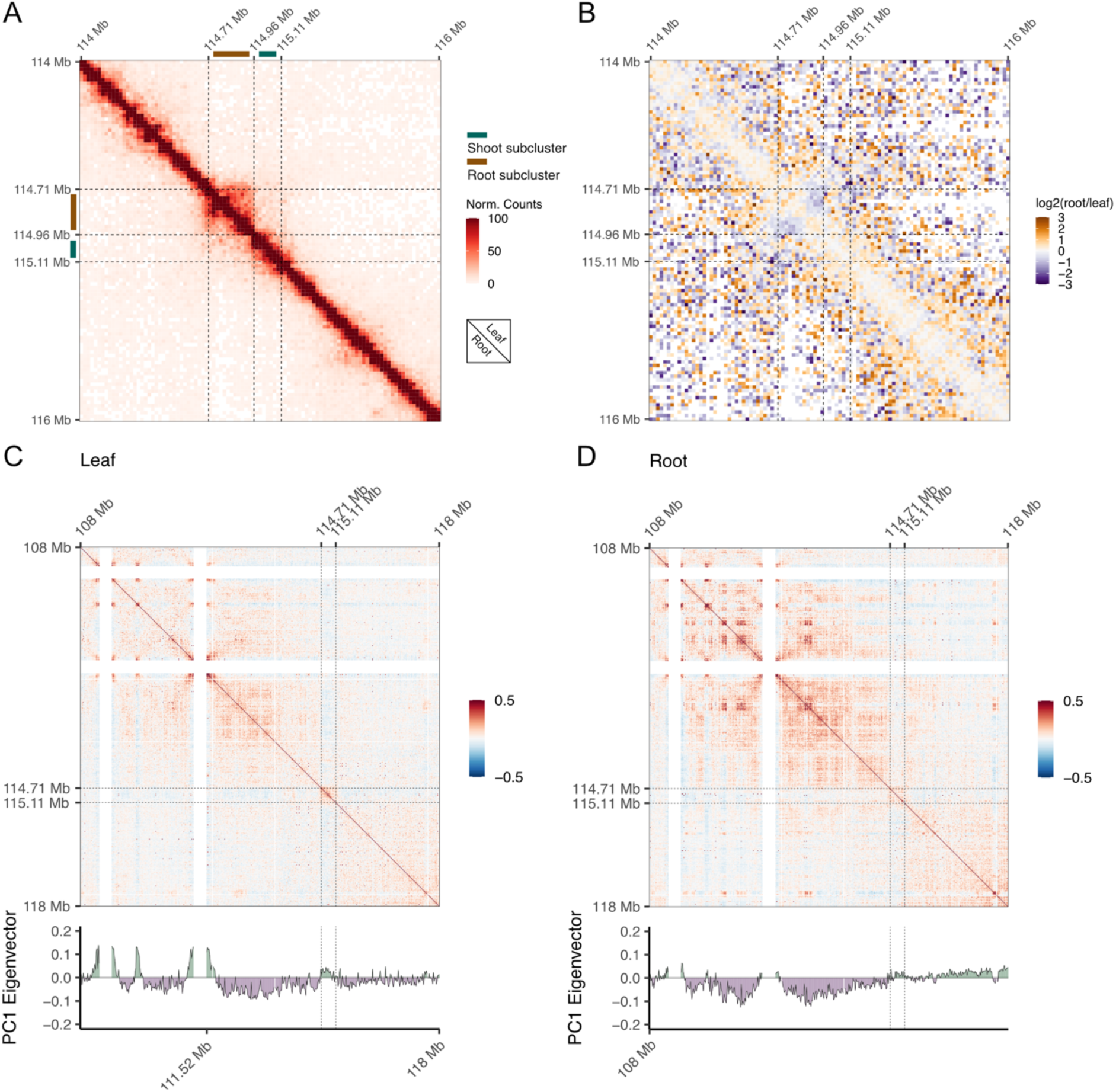
Higher-order chromatin configuration and chromatin-chromatin interactions at the withanolide BGC locus. **A)** Normalised Hi-C contact matrices at 20-kb resolution covering the locus of the withanolide BGC of leaf (above the diagonal) and root (below the diagonal). **B)** Binary logarithm (log2) of the ratio between the root and shoot normalised Hi-C contact matrices at 20-kb resolution. Positive and negative values indicate more interaction frequency in root and leaf tissue, respectively. **C)** and **D)** Pearson’s correlation matrices of chromosome 1 right arm and sub-A/B compartment annotation in **C)** leaf and **D)** root. In the four panels, vertical and horizontal dashed black lines indicate the withanolide BGC locus.

We noticed that the entire withanolide BGC locus displayed very few interactions with the neighbouring euchromatic region, visible as a banded area in the Hi-C map, which was more pronounced in the shoot than the root (**Supplementary Figure 7D,E**). This led us to hypothesise that the local chromatin state of the whole BGC might differ from that of the neighbouring regions. Global euchromatic regions (A compartments) and heterochromatic regions (B compartments) can be further classified into local A/B compartments, i.e, local regions within the A or B compartments that can be either euchromatic or heterochromatic at a smaller scale and that correlate with activating or inactivating epigenetic marks (Dong et al., 2017). Consequently, we annotated local A/B compartments within the euchromatic region containing the cluster (**Figure 3C,D**). In leaf tissue, the BGC clearly stood out from the neighbouring chromatin, suggesting that it preferentially interacted with heterochromatin (**Figure 3C**). This was less prominent in root tissue, where the cluster seemed to interact with the downstream euchromatic regions (**Figure 3D**). Collectively, the ATAC-seq and Hi-C results suggested that the local 3D organisation of the chromatin contributes to the regulation of the tissue-specific expression of each subcluster.

### The tissue-specific gene expression of each subcluster is associated with higher levels of asymmetric DNA methylation

Local epigenomic configurations are emerging as a key regulator of plant specialised metabolism (Conneely et al. 2022; Méteignier et al. 2022). The tissue-specific chromatin organisation at the withanolide BGC locus, revealed by our ATAC-seq and Hi-C data, led us to hypothesise that distinct epigenetic states might be contributing to the regulation of the bipartite sub-cluster-architecture of the withanolide BGC. Cytosine methylation at the 5th carbon (5mC, or often more generally referred to as DNA methylation) is an important epigenetic mark involved not only in gene expression regulation and the silencing of transposable elements but also in modulating chromatin interactions (Zhang et al. 2018). Moreover, recent studies showed that the expression of biosynthesis genes in several specialised metabolism pathways are negatively correlated with DNA methylation (Li et al. 2019; de Bernonville et al. 2020; Kim et al. 2020). We therefore hypothesised that the withanolide BGC sub-cluster architecture might be associated with distinct DNA methylation patterns. We performed whole-genome bisulfite sequencing (WGBS) of shoot and root tissue of 2-week-old seedlings, from the exact same biological samples as for the RNA-seq and untargeted metabolomic analyses. We sequenced three biological replicates per tissue, with a mean sequencing depth of 23-40X and bisulfite conversion rates >99.3% in every sample (**Supplementary Dataset 4**; **Supplementary Figures 8 and 9**). We called differentially methylated regions (DMRs) between shoot and root samples (Hüther et al. 2022). In a principal component analysis (PCA) of mean methylation rates in context-specific DMRs, samples clustered by tissue in all three contexts (**Supplementary Figure 8A**). In total, we identified 322 CG-, 183 CHG- and 9,522 CHH-context-specific DMRs. The withanolide BGC region contained 0 CG-, 1 CHG-, and 20 CHH-DMRs (**Figure 4A**). Contrary to our expectation, in each sub-cluster hypermethylated CHH-DMRs correlated with the active/expressed state (**Figure 4A**). Of the 20 CHH-DMRs in the withanolide BGC, 4 and 2 were located in the putative promoters (2 kb upstream) of RESC and SESC genes, respectively. Another CHH-DMR was located in the intron of the RESC gene *CYP87G1;* contrary to the trend, this gene was hyper-methylated in shoot tissue. The remaining 11 CHH-DMRs were located in intergenic regions.

**Figure 4.**
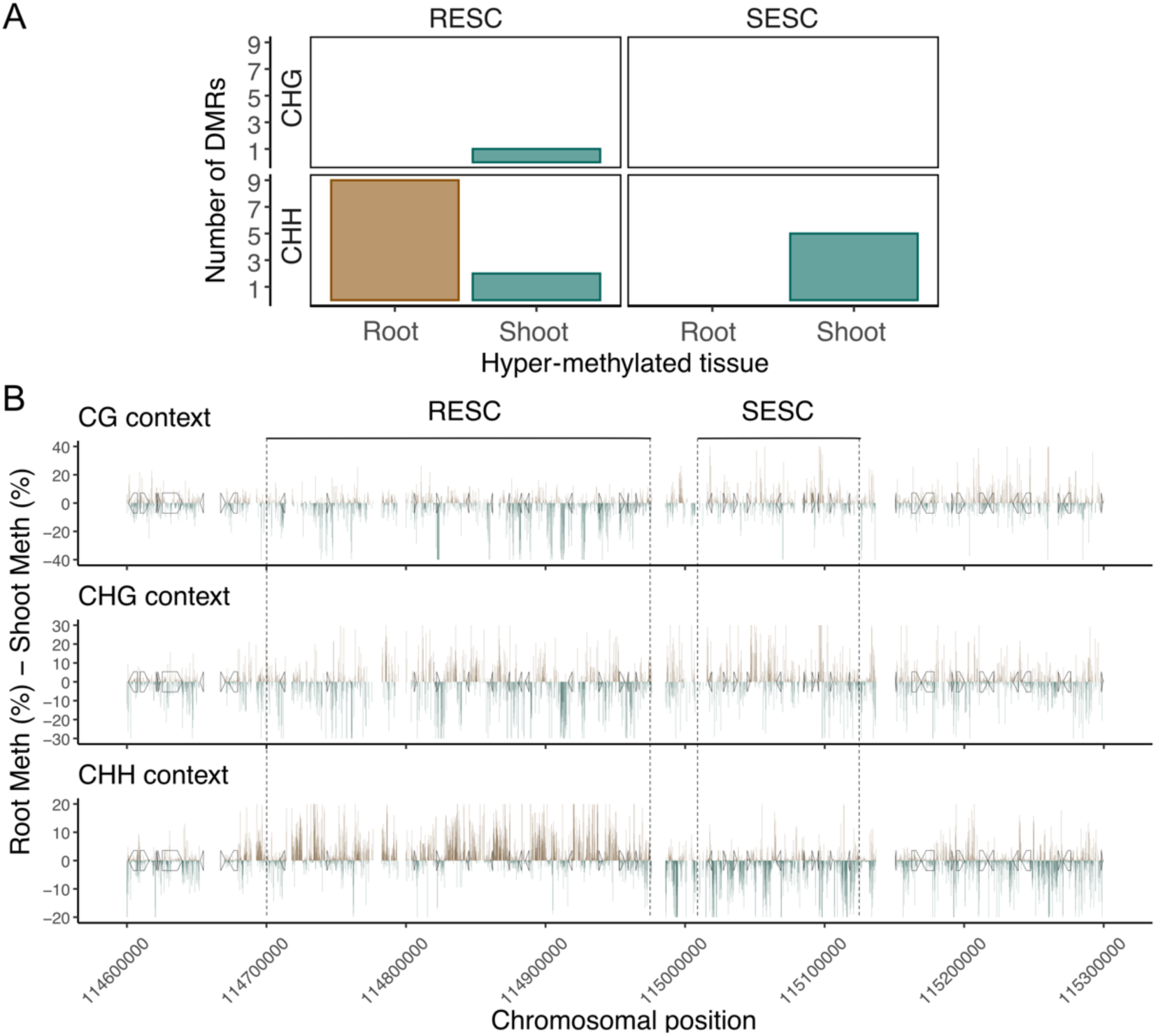
DNA methylation at the withanolide BGC locus in *P. grisea*. **A)** Number of differentially methylated regions (DMRs) identified within the RESC and SESC, respectively, in the comparison of root vs. shoot tissue. **B)** Differential DNA methylation between root and shoot tissue along the BGC and its flanking regions. Average methylation was calculated in non-overlapping 100 bp windows in root and shoot for each sequence context separately; windows that did not have above-minimum coverage information for both tissues were removed. The data show the per-context difference between root and shoot methylation for each window; positive values (in brown) represent higher methylation in root; negative values (in green) represent higher methylation in shoot tissue.

At a global level, the RESC also exhibited considerably higher CHH methylation in root tissue, where that sub-cluster is expressed, compared to shoot, in line with the DMR hypermethylation (**Figure 4B**). The SESC displayed the opposite trend, i.e., CHH methylation was highest in shoot tissue. CHH methylation differences primarily occurred in intergenic regions. While we did not observe a noticeable pattern for CHG methylation, CG methylation followed the opposite, albeit less pronounced trend than CHH methylation: in the RESC, CG methylation was negatively correlated with expression in the respective tissue. In sum, our data showed a marked shift in tissue-specific DNA methylation occurring along the genomic region that separates the two sub-clusters.

### The withanolide BGC in Solanaceae underwent lineage-specific gene duplications

Withanolides are widespread across *Solanaceae* (Misico et al., 2011). While they are absent in most *Solanum* species, they have been isolated, among others, from the late-diverging *Physalideae* tribe, as well as from early-diverging *Petunia* (in the form of petuniasteroids), and are present in e.g. species from the genera *Datura*, *Nicandra* or *Lycium* (Hansel et al. 1975; Elliger et al. 1989; Misico et al. 2011; Huang et al. 2020; Pan et al. 2024).

Given the ubiquitous presence of withanolide-related metabolites in several *Solanaceae* genera, we asked how conserved the withanolide BGC was across the *Solanaceae*, and whether the tandem duplication and sub-functionalization was a feature commonly associated with this cluster. We mined the available *Solanaceae* genome assemblies, including species and genera known to produce withanolides (**Supplementary Table 2**). Using Orthofinder (Emms and Kelly 2019), we defined orthogroups; within the same orthogroup we searched for the orthologs most similar to the genes in the *P. grisea* withanolide BGC. Among the list of candidate orthologs in each surveyed species, we further confirmed which of these were physically clustered in the genome.

We identified orthologs of *24ISO* and of several of the *P. grisea* withanolide BGC genes located in what appeared to be BGCs across most of the tribes included in our study. Genes encoding for 24ISO orthologs and associated gene clusters were markedly absent in many *Solanum* species, particularly in tomato- and potato-related species (*S. lycopersicum*, *S. pennelli*, *S. stenotomum*, *S. verrucosum*, *S. septemlobum*, *S. lyratum*, *S. septemlobum*, *S. seaforthianum* and *S. americanum*). In turn, we found a cluster in eggplant-related and several other *Solanum* species (*S. aethiopicum*, *S. macrocarpon*, *S. torvum*, *S. atropurpureum*, *S. viarum*, *S. wrightii*), but withanolides have not yet been reported in any of these (**Figure 5A**; **Supplementary Figure 10**). Conversely, the genomes of all species with reported withanolide production (or of their close relatives) contained at least one *24ISO* copy located in a putative withanolide BGC. Synteny analysis of the gene cluster itself as well as of the flanking chromosomal environment revealed that both were largely syntenic, indicating that the withanolide BGC in *P. grisea* originated from an early common ancestral cluster that existed already before the branching off of *Petunia* (**Figure 5B**). The cluster appears to have been lost in several tribes and species, but conserved in many others.

**Figure 5.**
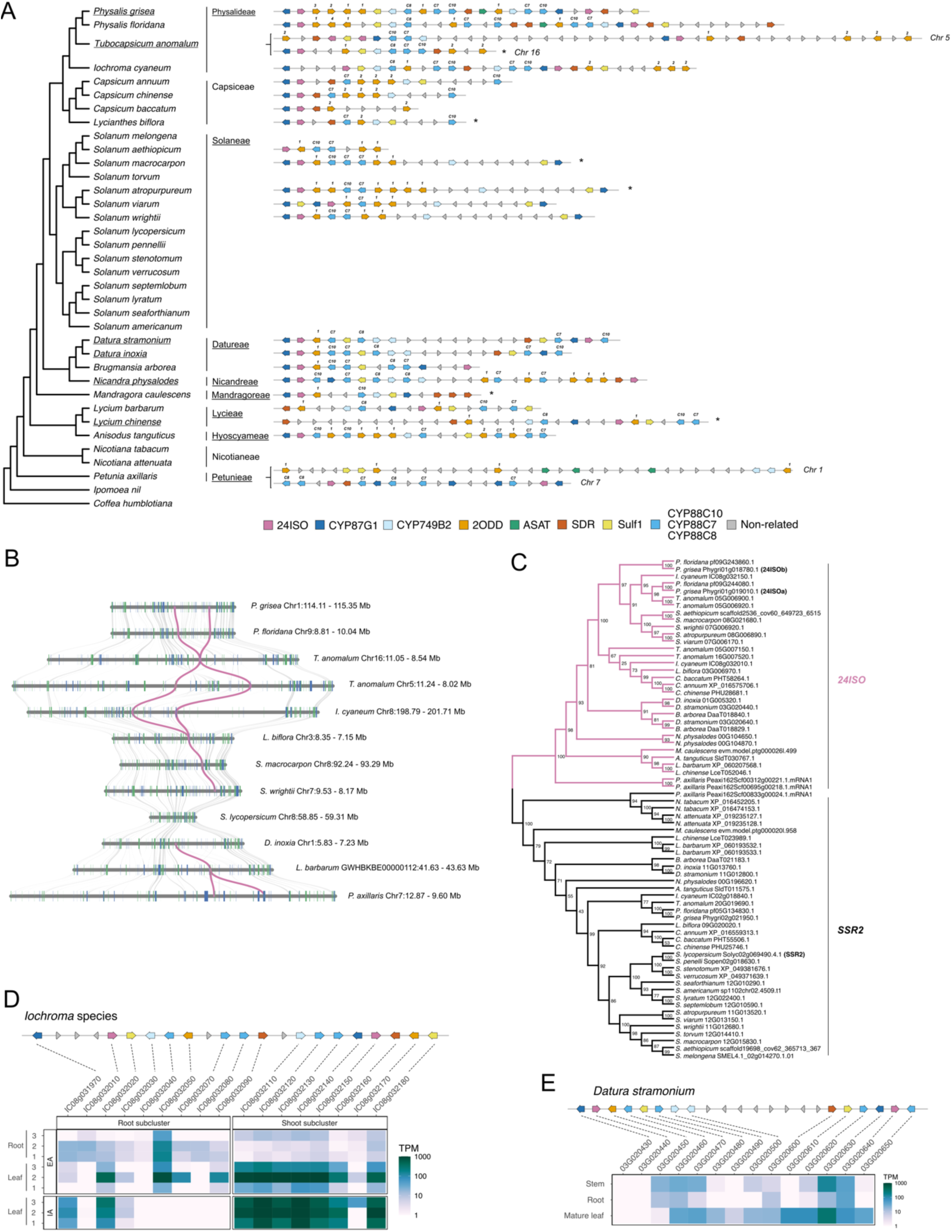
(next page). An ancestral withanolides biosynthetic gene cluster in *Solanaceae* underwent several lineage-specific gene duplications. A) Phylogenetic relationship between the different *Solanaceae* species included in the analysis (outgroups *Ipomoea nil* and *Coffea humblotiana*) and their respective withanolide BGC-like loci. Underlined species and/or tribes are those in which withanolide-related metabolites have been detected (**Supplementary Table 2**). An asterisk at the end of cluster indicates that this cluster has been inverted for representation purposes. Numbers above the *CYP88C* genes indicate the respective subfamily; numbers above 2ODD genes indicate the respective orthogroup that they belong to. **B)** Microsynteny between the genomic region containing the withanloide BGC in *P. grisea* and different representatives of the *Solanaceae* tribes included in this study. Coloured lines indicate *24ISO* orthologs. Grey lines connect other orthologs. **C)** Amino-acid-sequence maximum likelihood tree with the best-fit model selected by ModelFinder represented as a cladogram of the orthogroup containing the Pg24ISO. Numbers indicate ultra-fast bootstrap values. **D)** and **E)** Heatmap showing the expression pattern of each gene within the cluster across different tissues in the *Iochroma* species *Eriolarynx australis* (EA) and *Iochroma arborescens* (IA) **(D)** and in *Datura stramonium* (E). Expression levels are presented as transcripts per million (TPM).

The tandem duplication that resulted in two paralogs of *24ISO* and seems to have been at the origin of cluster sub-functionalization was apparent throughout the *Physalideae* tribe (including the genera *Physalis*, *Tubocapsicu*m, and *Iochroma*). To test whether this conserved BGC architecture was accompanied with conserved tissue-specific expression, we generated RNA-seq data from two *Physalideae* species that we could sample in the botanical garden in Munich, Germany. From the tree-like *Iochroma arborescens*, we could sample only leaf tissue; from the shrub *Eriolarynx australis* (also frequently listed as *I. australe*), we sampled leaf and root tissue (**Supplementary Dataset 1**). Our data showed that in leaves, the equivalent of the *P. grisea* SESC, i.e., the sub-cluster located towards the proximal end of the whole BGC, higher expression than its RESC equivalent (**Figure 5D**). Expression of the latter, on the other hand, was weaker in general but appeared to be slightly stronger in root tissue (only measurable for *E. australis*) (**Figure 5D**). We observed the same pattern in the SESC from *Tubocapsicum anomalum* in publicly available RNA-seq from young leaves (**Supplementary Figure 11**). Altogether, this indicates that the tandem duplication and sub- functionalization of the cluster is common to species of the *Physalidea* clade.

Our phylogenomic analysis also revealed several tribes with apparent cluster duplication, namely *Datureae*, *Nicandreae*, and *Petunieae*, usually accompanied by the core duplication of the *24ISO*, *CYP87G1* and the *CYP88C7* (**Figure 5A**). The amino-acid-sequence-based phylogenetic analysis of all the genes within the *P. grisea* withanolide BGC suggested that, in these tribes, these duplications were tribe-specific and unrelated to the duplication of the sub- cluster architecture in the *Physalideae* (**Figure 5C; Supplementary Figures 12-20**). To further test this hypothesis, we retrieved publicly available transcriptomic data for stem, root, and leaf tissue of *Datura stramonium*. In contrast to the *Physalideae* species, the cluster in *D. stramonium* did not show tissue-specific expression of different sub-clusters (**Figure 5E**). In line with the different cluster architecture in comparison to the BGC in *Physalideae*, this suggested an evolutionarily independent duplication event that did not result in sub- functionalization.

## Discussion

Next to frequent biosynthetic operons in bacteria, the organisation of collinear genes in BGCs has long been considered to be exclusive to fungi and plants; recent evidence suggests that they also exist in the animal kingdom (Scesa et al., 2022). BGCs likely ensure the co-inheritance of genes encoding for the subsequent steps in a biosynthetic cascade, which would safeguard against the accumulation of potentially toxic intermediates (Nützmann and Osbourn 2015). BGCs have also been proposed to facilitate the formation of metabolons, i.e., temporary complexes composed of the sequential enzymes of a biosynthetic process (Nützmann and Osbourn 2015). While these two aspects of BGCs remain hypothetical, it is certain that BGCs coordinate gene expression to ensure co-expression of the biosynthetic genes across tissues and treatments and hence the correct spatio-temporal production of the metabolites (Nützmann and Osbourn 2015). However, the mechanisms and temporal events that lead to the formation and diversification of BGCs remain unresolved (Polturak et al. 2022; Smit and Lichman 2022). Similarly, our understanding of BGC transcriptional regulation in plants is limited.

Here, we have identified a BGC underlying withanolide production in *P. grisea*. We show how a partial duplication resulted in the formation of two tandemly arranged sub-clusters, which are expressed in different tissues and appear to contribute to the production of tissue-specific withanolide compounds. Multi-layer epigenomic analysis revealed that the two sub-clusters are isolated from each other at the chromatin level, and that the tissue-specific expression is regulated epigenetically.

### An ancestral withanolide BGC with lineage-specific diversification in Solanaceae

By anchoring our search for a withanolide BGC on the *24ISO* orthologs in *P. grisea*, we identified a BGC-like arrangement of 22 genes, including several encoding CYP450s from different families and subfamilies, as well as six *2-ODD* genes. In terms of size (close to 400 kb) and gene content, this makes the withanolide BGC one of the largest plant BGC identified to date, comparable to the benzylisoquinoline alkaloid BGC in opium poppy (*Papaver somniferum*; 500 kb, 15 genes) (Guo et al. 2018). VIGS-mediated suppression confirmed the contribution of several genes to withanolide production (**Figure 1**). Our comparative phylogenomic and transcriptomic analyses showed that the withanolide BGC architecture and the division into two distinctly expressed sub-clusters are conserved across the *Physalideae*. The identification of orthologous clusters and independent duplication events across the *Solanaceae* family suggest the existence of an early ancestral cluster. We detected signals of at least two independent duplication events, one in the *Physalideae* clade, and one in the

### *Datureae*/*Nicandreae*/*Petunieae* clade, which resulted in different architectures and functionalities

In contrast, our analyses indicate the loss of the withanolide BGC in many species of the genus *Solanum*, particularly in those more closely related to tomato and potato. This is noteworthy, since it has been postulated that the biosynthesis of steroidal glycoalkaloids (prominent in tomato and potato and presumably in many related species) and that of withanolides, which branch off the same precursors, usually do not co-occur in the same species. Few exceptions to this have been reported to date, including *D. stramonium*, which produces both steroidal glycoalkaloids and diverse withanolides (Pan et al. 2024), and in which we identified an orthologous withanolide BGC (**Figure 1**). Although we found putative withanolide BGCs in several *Solanum* species, evidence of withanolide production in these species remains to be confirmed in future studies. Conversely, withanolides have been detected outside the *Solanaceae* family, and it will be interesting to investigate the genetic architecture underlying withanolide production in those species.

### A sub-functionalisation that results in tissue-specific activity

Our transcriptomic transect across different tissues, developmental stages and treatments revealed that the proximal and distal parts of the withanolide BGC in *P. grisea* are expressed tissue-specifically. In particular, the proximal set of 14 genes (the RESC) showed a strictly root-specific expression that was also maintained upon stimulating elicitors such as MeJA. The expression of the eight distal genes (the SESC) was most prominent in the aerial tissues, albeit less tightly restricted than the RESC. In line with this finding, we detected tissue-specific accumulation of different withanolide compounds. This suggests that the expression of the respective paralogs in either root or shoot might lead to the production of slightly different withanolide compounds and raises the question about the physiological and ecological function of this compartmentalisation. Since withanolides play a role in organismic interactions and act as antibiotics as well as in the repelling of herbivores (Huang et al. 2020), one can speculate that root-specific and shoot-specific withanolides could play respective roles in the defence against niche-specific microbes, nematodes, or insects. In future, the genetic and biochemical characterization of the two-sub-clusters will help understand their role in the chemical ecology of *P. grisea* and its relatives.

In line with the differential transcriptional activity in roots and shoots, our ATAC-seq data revealed regions of increased local chromatin accessibility within the RESC and SESC, respectively. Recent studies in the medicinal plants *Artemisia annua* and *Catharanthus roseus* have shown that tissue- and even cell-type-specific local chromatin accessibility correlate with BGC expression and the biosynthesis of artemisinin and monoterpene indole alkaloids, respectively (Zhou et al. 2021; Li et al. 2023). Such tissue-specific local chromatin accessibility allows TFs and other DNA-binding proteins to bind and to specifically activate gene expression. The next step in elucidating withanolide BGC expression will therefore be to identify the respective TFs and their binding sites. Our motif search of accessible regions returned bHLH and MYB TF binding motifs in root, and C2H2 zinc-finger binding motifs in shoot tissue (**Supplementary Table 3**), which will serve as an anchor for the future search of regulating TFs. An alternative approach is to search for TFs that are co-expressed along with either the RESC or SESC. Using our RNA-seq data across tissues and treatments, we identified, among others, two basic helix-loop-helix (bHLH) TFs in the corresponding expression modules. Members of this TF family are generally known to regulate specialised pathways through jasmonate (JA) signalling (Goossens et al. 2017). *Phygri08g015540* was co-expressed with the SESC in module M1. Interestingly, its closest homolog in tomato (*Solyc01g096370*) has been shown to be involved in the regulation of steroidal glycoalkaloids (Li et al. 2020). In module M2, which contained the RESC genes, we identified *Phygri08g006590*, whose closest homolog in Arabidopsis (bHLH20/NAI1; AT2G22770) is involved in glucosinolate accumulation and defence (Gao and Dubos 2024). Follow-up genetic analysis of these and other candidate TFs, including e.g. ethylene response factors (ERFs), which have been shown to regulate terpenoid biosynthesis across various plant lineages (Shoji et al. 2023), is necessary to elucidate the regulatory mechanisms that control sub- cluster expression.

### The sub-functionalisation of the withanolide BGC is maintained at the chromatin level

The withanolide BGC in *P. grisea* combines two features that have been reported for other plant BGCs individually, but never in combination: a defined organisation at the chromatin level, and a bipartite architecture. Recent studies have shown that plant BGCs can be organised in TADs or TAD-like structures, i.e., in regions with extensive 3D chromatin-chromatin interactions, and that the configuration of the TAD influences the (co-)expression of the genes located within the BGC. For example, in leaves of *A. thaliana*, the thalianol cluster is contained in a condensed B compartment and transcriptionally inactive, while in roots it is located in a relaxed-chromatin A compartment and transcriptionally active (Liu et al. 2020). In opium poppy, the BGC encoding the biosynthesis of benzylisoquinoline alkaloids also forms a TAD (Guo et al. 2018). Liu et al. recently reported on a *p*-menthane BGC in a *Lamiaceae*, *Schizonepeta tenuifolia*, that underwent an inverted duplication to form a bipartite cluster (Liu et al. 2023). Another *Lamiaceae* cluster, the miltiradiene BGC, exhibits gene duplications and even displays tissue-specific expression differing along different parts of the cluster (Bryson et al. 2023). The withanolide BGC described in our study exemplifies how a duplication of parts of a BGC can result in the formation of two tandemly arranged sub-clusters that end up being isolated from each other at the chromatin level: the two sub-clusters are arranged in two separate TAD-like domains. Their condensation levels differed between tissues and negatively correlated with expression; especially the RESC showed more intense interactions in leaves than in roots. Since the RESC was not expressed in the shoot (and *vice versa* the SESC was only lowly expressed in roots), the Hi-C, along with the ATAC-seq data, suggest that both sub-clusters adopt a more compacted conformation in the tissue in which their expression is suppressed. Such a compaction likely restricts the accessibility for TFs and the primary transcriptional machinery. These observations are in line with the patterns observed for the thalianol cluster in *A. thaliana* (Liu et al. 2020; Conneely et al. 2022; Méteignier et al. 2023).

Unexpectedly, asymmetric CHH methylation was positively correlated with gene expression in both sub-clusters. While CHH methylation is generally associated with transcriptional suppression, it should be noted that this relies mostly on data from *A. thaliana* and several grass crops, and that our understanding of the epigenome in *Solanaceae* is rather limited. To our knowledge, this is the first observation of differential DNA methylation correlating with BGC activity. Further investigations into the chromatin landscape of *P. grisea* and the withanolide BGC are necessary, e.g. on the role of post-translational histone modifications and variants, which have been shown to be associated with structural organisation of the thalianol BGC (Liu et al. 2020), as well as on the potential impact of the gene-depleted 50-kb spacer sequence that separates the two sub-clusters and that features a prominent accessible chromatin region in its centre (**Figure 2B**).

## Conclusion

In summary, our study highlights how, based on chromatin- and epigenome-level organisation, eukaryotic BGCs can diversify and gain additional functionality. It moreover showcases that *P. grisea*, an emerging genetic model species in the *Solanaceae* family (Van Eck 2022; Dale et al. 2024), is amenable to further investigations into the biosynthesis of withanolides, a group of diverse and multi-functional plant specialised metabolites that are of interest in the context of plant protection and human health (Huang et al. 2020). We are aware that the tissue-specific analysis of the withanolide BGC presented here might yet mask a more refined structuring of withanolide biosynthesis, one that might be resolved in future by analyses at the single-cell level.

## Materials and Methods

### Plant material

Seeds for *P. grisea* were kindly provided by Sebastian Soyk (University of Lausanne), from the line sequenced for the *P. grisea* genome (He et al. 2023).

### Virus-induced gene silencing

Gene silencing was done using a tobacco rattle virus (TRV)-based VIGS system (Dinesh-Kumar et al. 2003) as described in (Knoch et al. 2018). Inserts were cloned into pTRV2 using XbaI and KpnI restriction sites and the resulting plasmids cloned into *Agrobacterium* GV3101. Fragments for genes to be silenced, except for Pg24ISO, were amplified from plasmids containing the corresponding *P. grisea* cDNA gene sequences. Primers used in this study are listed in **Supplementary Table 4**. The construct to silence Pg24ISO was synthesized by Azenta (Genewiz) based on the Ws24ISO construct described in Knoch et al (2018), with XbaI and KpnI restriction sites added. pTRV2 containing a *P. grisea* phytoene desaturase (PDS) fragment was used to verify VIGS and to determine time of harvest. pTRV2 containing a fragment of GFP was used as negative control. True leaves of 3-week-old *P. grisea* seedlings were infiltrated with *Agrobacterium* harboring pTRV1 and pTRV2 in a 1:1 ratio by syringe infiltration. Six plants were infiltrated per construct. Bleaching of PDS control plants was beginning about 3 weeks after infiltration and samples were harvested about 4 weeks after infiltration. The five top leaves from each plant were harvested and pooled for one sample, and aliquots of the same sample were used for qPCR and metabolomics analysis.

### Quantitative real-time PCR (qPCR)

RNA was extracted using RNeasy Plant Mini Kit (Qiagen) including on-column Dnase treatment. cDNA was synthesized using the RevertAid H Minus First Strand cDNA Synthesis Kit (Thermo Scientific). For qPCR 2 µl 100x diluted cDNA was used in 10 µl reaction volume. Real-time PCR was done with the QuantStudio5 (Applied Biosystems) system using FastStart Universal SYBR Green Master (Roche). Primers used in the analysis are listed in **Supplementary Table 4**. Expression levels in VIGS plants were calculated as 2^(-ΔΔCT)^ for each gene relative to Tubulin and GFP control plants. Five or six biological replicates were analyzed for each construct.

### Elicitor treatment

Seeds were sterilized for 15 min in 6% NaClO, washed 1x with 100% EtOH and 3x with water, and germinated on ½ Murashige & Skoog medium. Seedlings were grown horizontally in three rows on square plates in a Polyklima chamber at 24°C, 16 hours light (160 µmol), 8 hours dark cycle and 50% relative humidity for 2 weeks. For elicitor treatment, seedlings were transferred to Petri dishes (15 seedlings per dish, 4 replicates per condition) containing 30 mL water and left overnight on the bench. Water was removed and replaced with 30 mL elicitor solution (50 µM MeJa, 10 µM ABA, 1 µM Flg22, or mock); seedlings were incubated for 4 h before harvesting. Seedlings were dried with tissue paper; shoots and roots were separated with a razor blade. The tissue was flash-frozen in liquid nitrogen and ground to a fine powder.

### RNA-sequencing and transcriptomic analysis

Plants were grown in standard potting soil in the greenhouse. Different tissues were sampled as follows from 4 individuals: 1) young leaves, not fully expanded; 2) mature leaves, first fully expanded leaf; 3), petioles from mature leaves; 4) stems, top 10 cm; 5) calyx; 6) fruit, unripe, green stage. Seedlings for harvesting of two-week old roots and shoots were grown in sand at 24°C, 60 % relative humidity, and a 16 h light / 8 h dark cycle. Six to ten seedlings were pooled per sample, for a total of four biological replicates. Harvested tissue was flash-frozen in liquid nitrogen and ground to a fine powder. Samples were stored at -80°C until analysis and aliquots of the same sample were used for metabolomic and transcriptomic analysis.

RNA was extracted using the RNeasy Plant Mini Kit (Qiagen). In each set, each tissue/condition was replicated four times. After quality control, we discarded one replicate each for “fruit” and one for “young leaf” from the subsequent analyses (**Supplementary Figure 2**). We prepared polyA-RNA sequencing libraries from 400 ng of total RNA input using the NEBNext Ultra II RNA library preparation kit (New England Biolabs). Libraries were sequenced at IMGM (Martinsried, Germany) on a Novaseq instrument (Illumina) single-end 100 bp reads. Sequencing and mapping are summarised in **Supplementary Dataset 1**; quality control is shown in **Supplementary Figure 2**.

*Iochroma* samples were collected in the Botanical Garden in Munich, Germany. A pool of 2-3 young leaves from each plant were used for each replicate. Root samples could be harvested from only one plant; we sampled tissue from three different positions. *Iochroma* samples, RNA was extracted with Quick-RNA plant Miniprep Kit (Zymo Research).

For all the RNA-seq datasets, RNA-seq mapping and quantification was done using the nf-core pipeline rnaseq v3.9 (https://nf-co.re/rnaseq/3.9) implemented in Nextflow 21.10.3 (Di Tommaso et al. 2017; Ewels et al. 2020). Briefly, adapter and quality trimming was performed using TrimGalore v0.6.7 (Krueger et al., 2021), a wrapper tool around Cutadapt (v3.4) (Martin 2011) and FastQC (v0.12.1) (https://www.bioinformatics.babraham.ac.uk/projects/fastqc/). Reads were mapped to the *P. grisea* (He et al. 2023), *I. cyaneum* (Powell et al. 2022), *T. anomalum, L. biflora* and *D. stramonium* (Wu et al. 2023) genome assemblies using STAR v2.7.10a (Dobin et al. 2013) and quantified using salmon v1.5.2 (Patro et al. 2017). Feature counts were normalised by gene length and sequencing depth using transcripts per million (TPM), that were used to inspect the expression of the withanolide BGC locus. Differential gene expression was performed with the DESeq2 (v1.44.0) R package (Love et al. 2014) using raw feature counts. For the co-expression analysis, WGCNA implemented in the R package CEMiTool (v1.28.0) was used (Zhang and Horvath 2005; Langfelder and Horvath 2008; Russo et al. 2018). As input, the raw counts of *P. grisea* tissue and elicitors RNA-seq datasets were used after *vst* transformation (Love et al. 2014).

In the case of *T. anomalum, L. biflora* and *D. stramonium*, reads were obtained from the repository described in the original manuscript (Wu et al., 2023). At the time we mapped the first time the RNA-seq in *P. grisea* assembly, we noticed that several RESC genes (*CYP87G1b*, *2ODD4*, *CYP749B2b* and *CYP88C10*) were mis-annotated, probably due to the lack of expression of these genes in aerial tissue, used to annotate the genome in the original manuscript (Lemmon et al. 2018; He et al. 2023). Accordingly, we manually re-annotated those genes to a full CDS (re-annotation can be found in the manuscript repository).

### ATAC-seq data generation and analysis

Plants were grown in standard potting soil in the greenhouse. For each replicate, approximately ∼ 300 mg of young leaf and ∼ 1g of root of a pool of 2-3 3-week-old seedlings were harvested to isolate nuclei. The nuclei isolation protocol was based on Sikorskaite et al. (Sikorskaite et al. 2013) with slight modifications. Briefly, nuclei were extracted from the tissue on a petri dish on ice by chopping with a razor blade in nuclei isolation buffer (NIB: 10 mM MES-KOH pH 5.4, 10 mM NaCl, 200 mM sucrose, 0.1 mM spermine, 0.5 mM spermidine, 1 mM DTT and 1% BSA, 0.35% TritonX-100 - in the case of leaf tissue- or 0.75% for root tissue). The supernatant was filtered through a 40 μm strainer and the nuclei were pellet by centrifuging at 500 rcf for 5 min at 4°C. The pellet nuclei were resuspended in NIB, always using bored-open pipette tips, and filtered with a 20 μm strainer. The filtered nuclei were added on top of a NIB + 35% percoll solution and centrifuged at 500 rcf for 10 min at 4°C. The supernatant was removed and the nuclei resuspended in TAPS solution (10mM TAPS-NaOH, pH8.0; 5 mM MgCl2). We then dyed the nuclei with 1 μg/mL DAPI of 1 replicate of leaf and root. We estimated the number of nuclei in the root and leaf sample with a hemocytometer. We treated ∼50k nuclei with 2.5 μL of Tagment DNA enzyme (Illumina) in 25 μl of Tagment DNA Buffer in a final volume of 50 μL at 37°C for 60 min. The DNA was then purified using the Monarch® PCR & DNA Cleanup Kit (New England Biolabs #T1030). The tagmented DNA was amplified with the NEB II Ultra Q5 with 11 cycles using primers previously described in https://www.iaea.org/sites/default/files/warthmann-illumina-multiplexlibrary-protocol-2021.pdf. The DNA was recovered using AMPure beads. As an input sample, we extracted gDNA from a 3-week-old seedling that was subjected to the same tagmentation procedure and amplification. Libraries were sequenced at Novogene Germany on a NovaSeqX Plus platform (Illumina), using 2 x 150 bp paired-end reads.

ATAC-seq reads quality control and mapping was performed using the nf-core pipeline atacseq (v2.1.2) (https://nf-co.re/atacseq/2.1.2) implemented in Nextflow v23.10.0 (Di Tommaso et al. 2017; Ewels et al. 2020; Patel et al. 2023). Briefly, quality and adapter trimming of raw reads was performed using TrimGalore! v0.6.7 (https://github.com/FelixKrueger/TrimGalore). Read mapping was performed with bowtie2 (v2.4.4) (Langmead and Salzberg 2012). After mapping, duplicates were marked with Picard (v3.0.0) (https://broadinstitute.github.io/picard/). Subsequently, reads mapping to mitochondrial and chloroplast genomes or to blacklisted regions were removed using samtools (v1.17) (Danecek et al. 2021). Additionally, reads that were marked as duplicates, not marked as primary alignments, unmapped or mapping to multiple locations were also removed with samtools. Reads containing more than 4 mismatches, were soft-clipped or had an insert size > 2kb were removed with bamtools (v2.5.2) (Barnett et al. 2011). The alignment-level QC for the resulting filtered bam files were evaluated with picard. Normalised bigWig files scaled to 1 million mapped reads were generated using bedtools genomecov (v2.30.0) (Quinlan and Hall 2010) and bedGraphToBigWig (v445) (Kent et al. 2010).

The resulting filtered bam files from the pipeline were used to call narrow peaks with hmmratac (Tarbell and Liu 2019) implemented in MACS3 (3.0.2) (Zhang et al. 2008). For each tissue, we used all the replicates as input. Leaf and root accessible regions were filtered for a score >100. Bedtools intersect was used to get consensus narrow peaks of leaf and root tissues. The raw counts of consensus narrow peaks were used to compute the PCA with DESeq2. The Homer (c4.11) annotatePeaks and findMotifsGenome algorithms (Heinz et al. 2010) were used to annotate narrow peaks on each tissue and to find enriched motifs in the RESC (using root narrow peaks) and in the SECS (using leaf narrow peaks). Normalised bigWig files were used as input for deeptools (v3.2.0) (Ramírez et al. 2016) to compute metaplots over the annotated genes scaled to 3kb. Visualisation of the chromatin accessibility of the withanolide BGC locus was performed with igv (Robinson et al. 2011).

### Hi-C data generation and analysis

Leaf samples and root samples consisted of a pool of two and four seedlings, respectively. Approximately 0.5 g of fresh tissue was collected and fixed with 1% formaldehyde in MC buffer (10 mM potassium phosphate, pH 7.0; 50 mM NaCl; 0.1M sucrose; 0.01% Triton X-100) by applying a vacuum for 30 mins at room temperature. Formaldehyde was neutralised by washing the samples with MC buffer supplemented with 0.15 M glycine and applying a vacuum for 10 min at room temperature. Subsequently, the tissue was rinsed with fresh MC buffer twice for 5 min and further washed with water. Next, the tissue was snap-frozen and stored at -80 until library preparation.

Library preparation was started with tissue homogenization with ice-cold nuclei isolation buffer (20 mM HEPES, pH 8.0, 250 mM sucrose, 1 mM MgCl2, 5 mM KCl, 40% glycerol, 0.25% Triton X-100, 0.1 mM PMSF, 0.1% 2-mercaptoethanol); subsequently, the homogenate was filtered with double-layered miracloth (Millipore). After washing twice with nuclei isolation buffer and one round with 1× NEB Dpn II buffer (NEB), nuclei pellet was resuspended with 250 µl 0.5% SDS and split into five tubes. The subsequent steps followed essentially a published protocol (Wang et al. 2023). Nuclei penetration was then performed at 62 °C for 5min, followed by adding 145 µl water and 25 µl 10% Triton X-100 each tube to quench SDS. Subsequently, chromatin digestion was carried out at 37°C overnight with 50 U DpnII (NEB) in each tube, and stopped by incubating at 62°C for 20 min the next day. For sticky end filling and chromatin labelling, 1 µl of 10 mM dTTP, 1 µl of 10 mM dATP, 1 µl of 10 mM dGTP, 10 µl of 1 mM biotin- 14-dCTP, 29 µl water and 40 U Klenow fragment (Thermo Fisher) were added, and further incubated at 37°C for 2 h. After adding 663 µl water, 120 µl 10× blunt-end ligation buffer (300 mM Tris–HCl, 100 mM MgCl2, 100 mM DTT, 1 mM ATP, pH 7.8) and 40 U T4 DNA ligase (Thermo Fisher), proximity ligation was performed at room temperature for 4 h. Next, five tubes of nuclei pellet were combined by resuspending with 650 µl SDS buffer (50 mM Tris–HCl, 1% SDS, 10 mM EDTA, pH 8.0). After proteinase K (Thermo Fisher) treatment at 55 °C for 30 min, de-cross-linking was carried out by adding 30 μl 5 M NaCl and incubating at 65 °C overnight. The next day, following DNA recovery and RNA digestion with RNase A, 3 μg purified Hi-C DNA was sonicated in 130 µl TE buffer (10 mM Tris–HCl, 1 mM EDTA, pH 8.0) with a Q800R3 sonicator (QSONICA) by using the parameters: 25% amplitude, 15 s ON, 15 s OFF, pulse-on time for 4.5 min, to achieve fragment size shorter than 500 bp. Sonicated products were further purified with MinElute PCR purification kit (QIAGEN) and fragments larger than 300 bp were recovered with Ampure beads. Subsequently, the DNA was mixed with 0.5 μl 10 mM dTTP, 0.5 μl 10 mM dATP and 5 U T4 DNA polymerase (Thermo Fisher) in a 50 μl reaction volume and incubated at 20 °C for 30 min to remove biotin from unligated DNA. After DNA recovery with Ampure beads, end repair and adaptor ligation were performed following the manual of NEBNext Ultra II DNA Library Prep Kit (NEB). The ligated DNA was then affinity pulled-down with Dynabeads MyOne Streptavidin C1 beads (Invitrogen), and amplified with 13 PCR cycles.

Hi-C libraries were sequenced at the LAFUGA Gene Center Munich on a NextGen 500 instrument (Illumina) using 2 x 60 bp paired-end reads. Sequencing and mapping statistics are summarised in **Supplementary Dataset 3**; quality control is shown in **Supplementary Figure 7**.

Hi-C mapping was performed with the nf-core pipeline nf-core/hic (v2.1.0) (Servant et al. 2023) implemented in nextflow v24.04.2. Briefly, read quality control was performed with FastQC (v0.11.9). Hi-C data processing was done using HiC-Pro (Servant et al. 2015). HiC-pro employs bowtie2 (v2.4.4) to perform the mapping using a two-step strategy that rescues reads spanning the ligation site, following with the detection of valid interaction products and duplicates removal to finally generate the raw contact maps.

For all the samples and replicates, we randomly selected the same number of valid pairs (∼41 mio), in reference to the sample with the lowest number of valid pairs (leaf replicate 2). We used pre from juicer tools (1.9.9) (Durand et al. 2016b) to transform the raw contact maps from HiC-pro (.AllValidPairs) into .hic format. Downstream analyses were performed in the R environment (v4.4.1). Strawr (v0.0.92) (Durand et al. 2016a) was used to parse the .hic matrices into R with a resolution of 50,000 bp and normalised with the Knight and Ruiz (KR) method (Knight and Ruiz 2013). Quality control of replicates was performed applying Pearson correlation and representing the relationship of the samples with hierarchical clustering implemented in the R package pheatmap (v1.0.12).

After confirming the clustering of the replicates by tissue, we combined all the replicates of one tissue into one sample and re-run the nf-core/hic pipeline, to increase the number of valid pairs per sample. Pre was used to convert the raw contact maps into .hic format. Juicebox (Durand et al., 2016) was used to visually explore the genome-wide contact matrix. Hi-C matrices were parsed into R using strawr. The withanolide BGC locus was explored with KR normalised Hi-C matrices at 20-kb resolution. To compute the PCA for the A/B and sub-A/B compartment annotation, we calculated Pearson’s correlation of the KR normalised observed/expected matrices at 500-kb and 20-kb resolution.

### Whole-genome bisulfite sequencing and DNA methylation analysis

The identical samples as transcriptomic and metabolomic analyses were used. Genomic DNA (gDNA) was extracted using the DNeasy Plant Mini Kit (Qiagen). Libraries were prepared from 100 ng input gDNA as described in (10.1371/journal.pgen.1010452) using the NEBNext Ultra II DNA library preparation kit (New England Biolabs), in combination with the EZ-96 DNA Methylation-Gold MagPrep kit (Zymo Research) for bisulfite conversion. We used 0.01% λ- phage DNA for bisulfite conversion control. Libraries were sequenced at Novogene (UK) on a NovaSeqX platform (Illumina) with 2 x 150 bp reads. Sequencing and mapping statistics are summarised in **Supplementary Dataset 4**; quality control is shown in **Supplementary Figures 8 and 9**.

WGBS-seq reads quality control and mapping was performed using the nf-core pipeline methylseq (2.6.0) implemented in Nextflow v23.10.0 (Di Tommaso et al. 2017; Ewels et al. 2020, 2024), following the bwa-meth (v0.2.2) (Pedersen et al. 2014) and MethylDackel (v0.6.0) (https://github.com/dpryan79/MethylDackel) workflow. Briefly, the first three nucleotides from the 5’ end of read 1 and 2 were trimmed. Raw data quality control was performed with FastQC (v0.12.1) and adapter sequence trimming with cutadapt (3.4) implemented in Trimgalore (v0.6.7). After adapter trimming, the first three bases from the 3’ end of read 1 and 2 were also trimmed. Reads were aligned with bwa-meth (0.2.2). Duplicates were then marked with picard (v3.0.0). The methylation calls were extracted with MethylDackel (v0.6.0) with a minimum depth of 5 independent reads for MethylDackel to report a methylation call. Alignment quality control (estimation of genome coverage, insert sizes of mapped reads and GC content distribution) was performed with qualimap (v2.2.2- dev) (Okonechnikov et al. 2016). Alignment statistics were evaluated with samtools (v1.17).

MethylScore (v0.2) (Hüther et al. 2022) implemented in Netflow version v21.10.6 was used to call differentially methylated regions between all samples with default parameters. Non-conversion rate was calculated for the λ-phage genome as the total of reads supporting methylation for a methylated cytosine divided by the total number of reads. The per-site methylation frequency was calculated based on (Schultz et al. 2012) as the number of reads reporting a methylated base divided by the total reads mapping to that base. To calculate the relative frequency of methylation rates, sites with a coverage ≥10 were considered. Bedtools subcommands map and makewindows (v2.30.0) were used to calculate the mean site methylation in windows along the genome.

### Metabolite analysis

Metabolite extraction was performed as described in (Salem et al. 2020) with some minor modifications. Briefly, 1 ml of pre-cooled (−20°C) methyl tert-butyl ether:methanol (3:1 v/v) was added to homogenised tissues. After, tubes were thoroughly vortexed for 1 min and then incubated on an orbital shaker (1000 rpm) for 15 min at 4°C followed by sonication for 15 min. For phase separation, a volume of 500 μl of water:methanol (3:1 v/v) was added to each tube and the samples were again thoroughly vortexed for 1 min. After that, the samples were centrifuged at 13,300 g for 5 min. The dried polar aliquots were resuspended in water:methanol (1:1 v/v) and used for secondary metabolites analysis.

Secondary metabolites were profiled by the Waters Acquity ultra performance liquid chromatography (UPLC) system coupled to the Q Exactive Orbitrap mass detector according to the previously published protocol (Giavalisco et al. 2009). The UPLC system was equipped with a HSS T3 C18 reversed-phase column (100 × 2.1 mm i.d., 1.8-μm particle size; Waters, Milford, MA, USA) that was operated at a temperature of 40°C. The mobile phases consisted of 0.1% formic acid in water (Solvent A) and 0.1% formic acid in acetonitrile (Solvent B). The flow rate of the mobile phase was 400 μl/min, and 3 μl of sample was loaded per injection. The UPLC was connected to a Q Exactive Orbitrap focus (Thermo Fisher Scientific, Waltham, MA, USA) via a heated electrospray source (Thermo Fisher Scientific). The spectra were recorded using full scan and MS2 in both positive and negative ionisation detection, covering a mass range from m/z 100 to 1500. The resolution was set to 25 000, and the maximum scan time was set to 250 msec. The sheath gas was set to a value of 60, while the auxiliary gas was set to 35. The transfer capillary temperature was set to 150°C, while the heater temperature was adjusted to 300°C. The spray voltage was fixed at 3 kV, with a capillary voltage and a skimmer voltage of 25 and 15 V, respectively. Mass spectrometry spectra were recorded from minute 0 to 19 of the UPLC gradient. Molecular masses, retention time, and associated peak intensities were extracted from the raw files using refinerms (version 5.3; GeneData), and xcalibur software (Thermo Fisher Scientific). Metabolite identification and annotation were performed using standard compounds, data-dependent method (ddMS2) fragmentation, the literature, and metabolomics databases (Alseekh et al. 2021). To identify additional withanolide features, we searched the LOTUS database (Rutz et al. 2022) for withanolides and downloaded structure information for these. Using the exact mass, we identified withanolide structures in our dataset as those with a match in mass with less than 5 ppm difference to the exact mass of reported structures. Common leaf-specific withanolide features in the different tissues and in the VIGS untargeted metabolomic experiments were defined as features differing ≤0.02 in mass/charge ratio (m/z) and retention time.

### Withanolide BGC orthologs detection

OrthoFinder (v2.5.4) (Emms and Kelly, 2019, 2015) was used to infer orthogroups containing *P. grisea* withanolide BGC genes. Each orthogroup was aligned using MAFFT (v7.490) (Katoh et al. 2002; Katoh and Standley 2013). IQTREE (v2.1.4-beta) (Minh et al. 2020) was used to infer the maximum likelihood (ML) tree with the best-fit model selected by ModelFinder (Kalyaanamoorthy et al. 2017) with 1000 replicates of ultrafast bootstrapping (Hoang et al. 2018). Trees were represented with Dendroscope (v3.8.2) (**Supplementary Figures 12-20**). Genes within the same clade as *P. grisea* withanolide BGC genes were considered as potentially clustered homologs. The genomic location upstream and downstream of these homologs was further analyzed to confirm clustering.

Microsynteny analysis and figures were done with MCScan implemented in Python (v3.9) with JCVI utility libraries (v1.1.11) following the package workflow: https://github.com/tanghaibao/jcvi/wiki/MCscan-(Python-version). Briefly, protein FASTA files and GFF3 files of each species were used as input. When needed, the FASTA files were processed using Biopython’s SeqIO package (Cock et al. 2009). Pairwise ortholog and synteny blocks search was performed with default settings using a lift-over distance of 5, a c-score = 0.99 and a maximum of 3 iterations (6 in the case of *T. anomalum*). Multi-synteny blocks were constructed combining the syntenic blocks detected on every species used with respect to *P. grisea*.

### Data availability

The code used for all analyses will be made available upon final publication of the manuscript (https://github.com/spriego/Priego-Cubero_and_Knoch_et_al_Physalis). Sequencing data will be deposited in the ENA Short Read Archive (www.ebi.ac.uk/ena/) upon final publication under project no. PRJEB80835, submission no. ERA30854395.

## Author contributions

S.P.C., E.K., and C.B. conceived the study. S.P.C. carried out the genomic, epigenomic, transcriptomic, and phylogenomic analyses. E.K. carried out metabolomic analysis, and together with K.H.B and P.C. carried out molecular cloning and VIGS. Z.W. prepared Hi-C samples and analysed the data. S.A. prepared analytical samples and analysed metabolic profiles. A.R.F., C.L. and C.B. supervised the work. S.P.C. and C.B. wrote the manuscript with contributions from all authors.

## Acknowledgements

We would like to thank Niklas Schandry, Liza Rouyer and Peter Kerpan for critical reading of the manuscript and constructive feedback, and Zhihui Bao and Arlen Slaymaker for support in the laboratory work. We thank Simon Pfanzelt (Staatliche Naturwissenschaftliche Sammlungen Bayerns) for plant material of *E. australis* and *I. arborescens*, Sebastian Soyk (University of Lausanne) for providing *P. grisea* seeds, David Nelson (University of Tennessee) for naming the P450s, Norman Warthman (IAEA Vienna) for sharing the tagmentation protocol, Robert J. Schmitz (University of Georgia) for sharing the ATAC-seq protocol, and the LMU Biocenter Genome Centre for technical support. We acknowledge the Graduate School Life Science Munich (LSM) for their valuable support to S.P.C. This work was funded by the European Union’s Horizon 2020 research and innovation programme by the European Research Council (ERC), Grant Agreement No. 716823 ‘FEAR-SAP’ (C.B.); by the Deutsche Forschungsgemeinschaft (DFG) in the frame of TRR356 ‘Plant-Microbe’ (grant no. 491090170) and the project ‘Momidiverse’ (grant no. 457739273) to C.B.; and by the LMUexcellent programme by the Ludwig Maximilians Universität München (C.B., E.K.). For computational analyses, we used the BioHPC-Genomics compute cluster at the Leibniz Rechenzentrum (DFG; grant no. 450674345). S.A. and A.R.F. acknowledge the European Union’s Horizon 2020 research and innovation programme, project PlantaSYST (SGA-CSA No: 739582 under FPA No: 664620) and the BG05M2OP001-1.003-001-C01 project, financed by the European Regional Development Fund through the Bulgarian ‘Science and Education for Smart Growth’ Operational Programme,

## Supplementary Material

### Supplementary Figures

**Supplementary Figure 1.**
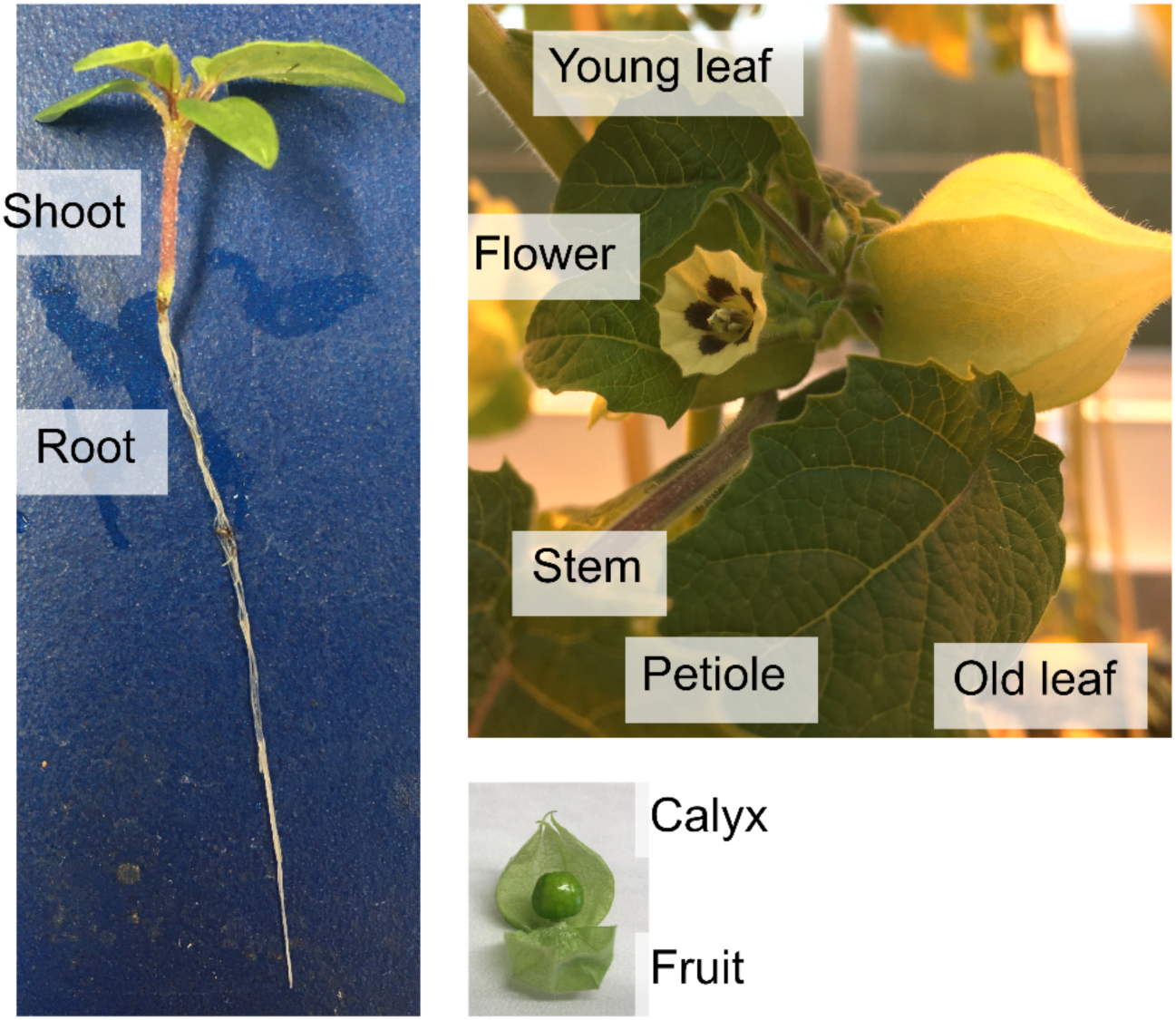
Overview of tissue selection for sequencing and metabolic analysis. Nine different tissues were selected for transcriptomic and metabolic analysis: root and shoot of 2-week-old seedlings (left); young leaf, mature leaf, petiole, stem, flower (open), calyx and fruit from adult plants (right). The same root and shoot samples were moreover used for WGBS library preparation.

**Supplementary Figure 2.**
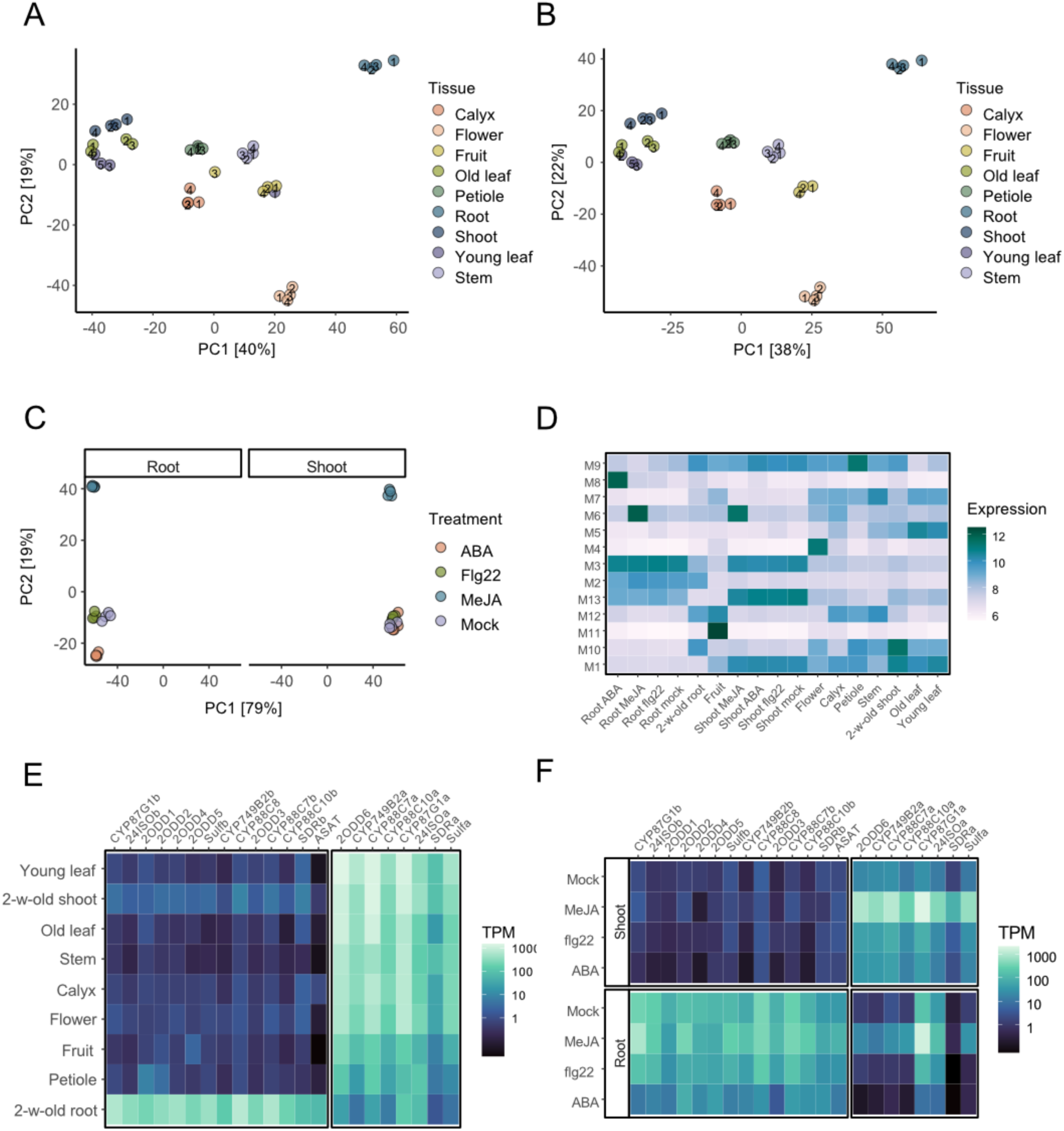
RNA-seq quality control and weighted gene correlation network analysis. Principal components analysis of **A)** ‘tissues’ RNA-seq before removing outliers, **B)** ‘tissues’ RNA-seq after removing two outlier samples, and **C)** ‘elicitors’ RNA-seq experiments using the top 500 genes with the highest variance after variance-stabilising transformation using the DESeq2 package (Love et al. 2014). **D)** Mean expression of the genes in each co-expression module (M1-M13) detected in the different samples (x-axis). The expression of each sample is the mean of four replicates (three replicates in the case of ‘Fruit’ and ‘Small leaf’, after outlier removal). The expression was normalised using the variance stabilising transformation. **E)** Heatmap showing the expression pattern of each gene in the withanolide BGC across different tissues. Expression levels are presented as transcripts per million (TPM). **F)** Heatmap showing expression in TPM of the genes within the cluster upon mock treatment and in response to various elicitors: methyl jasmonate (MeJA), flagellin 22 (Flg22), and abscisic acid (ABA).

**Supplementary Figure 3.**
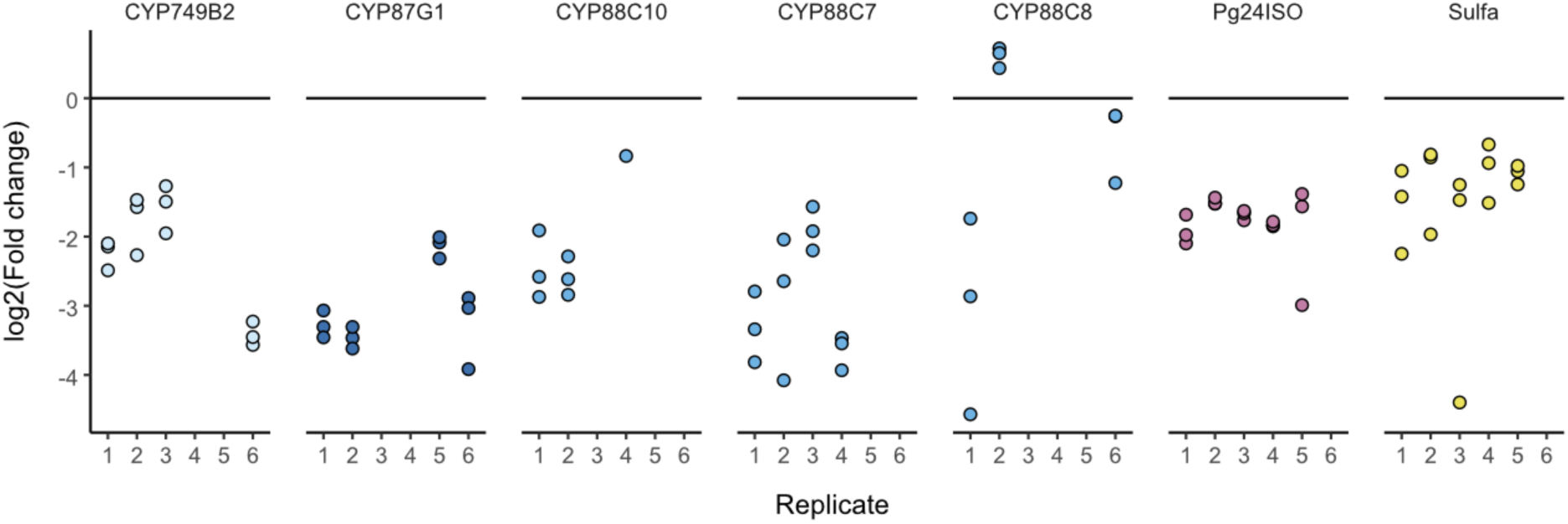
VIGS-induced silencing of biosynthetic cluster genes. qPCR was performed on VIGS-infiltrated *N. benthamiana* leaves in six biological replicates (1 to 6), with three technical replicates each, for the genes indicated at the top. Fold-changes are expressed in relation to an anti-GFP control (mock); all data were normalised against expression of a *TUBULIN* reference gene. Data is shown for those replicates that were selected, based on qPCR results, for the metabolic analysis in **Figure 1E**.

**Supplementary Figure 4.**
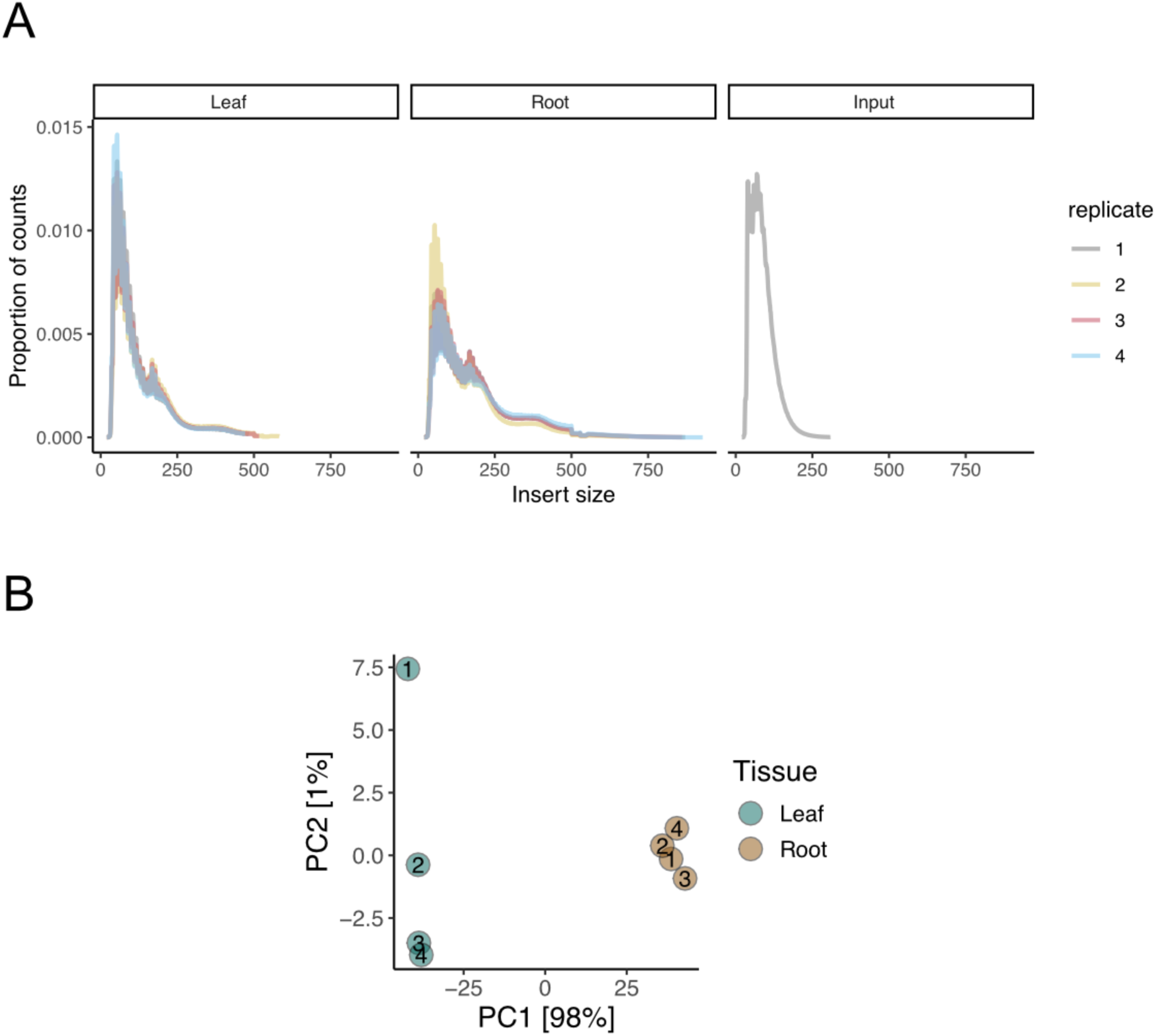
ATAC-seq quality control. **A)** Insert size (bp) distribution of ATAC-seq paired-end reads in leaf, root and input samples. **B)** Principal components analysis of ATAC-seq samples using the top 500 consensus ‘narrow peaks’ with the highest variance after variance-stabilising transformation using the DESeq2 package (Love et al. 2014).

**Supplementary Figure 5.**
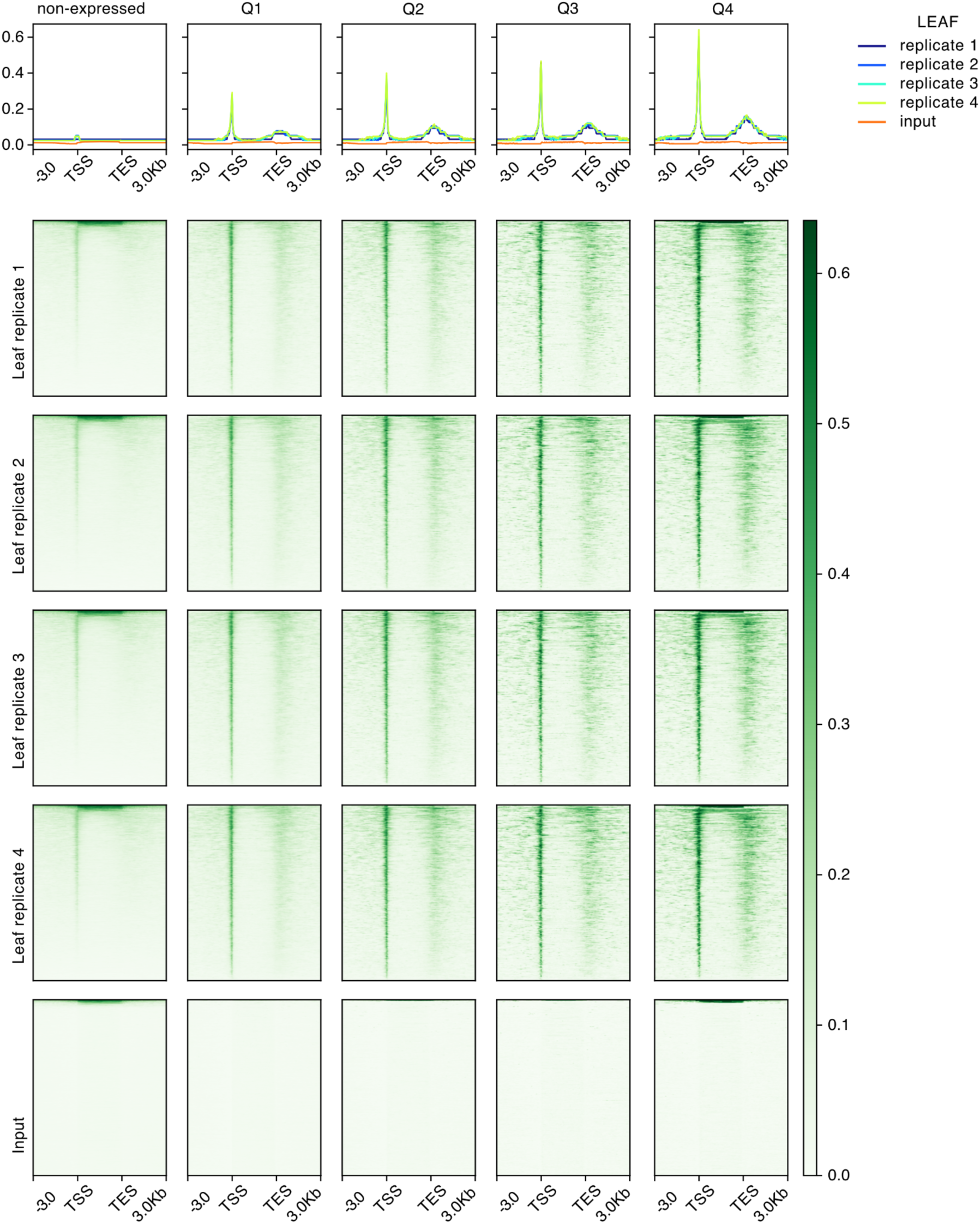
ATAC-seq metaplots of leaf samples. Metaplots and corresponding heatmaps of the genome-wide ATAC-seq signal (normalised by reads per million) of each leaf and input sample. Data was computed for non-expressed genes and for expressed genes divided into four quantiles of expression level (Q1=lowest to Q4=highest) (based on normalised gene expression in 2-week-old shoots). The plot represents 3 kb upstream (-3.0), transcription starting site (TSS), transcription end site (TES), and 3 kb downstream of each gene (3.0 kb). The gene body region was scaled to 3,000 bp. The profile plot represents the median of all the values for each group of genes.

**Supplementary Figure 6.**
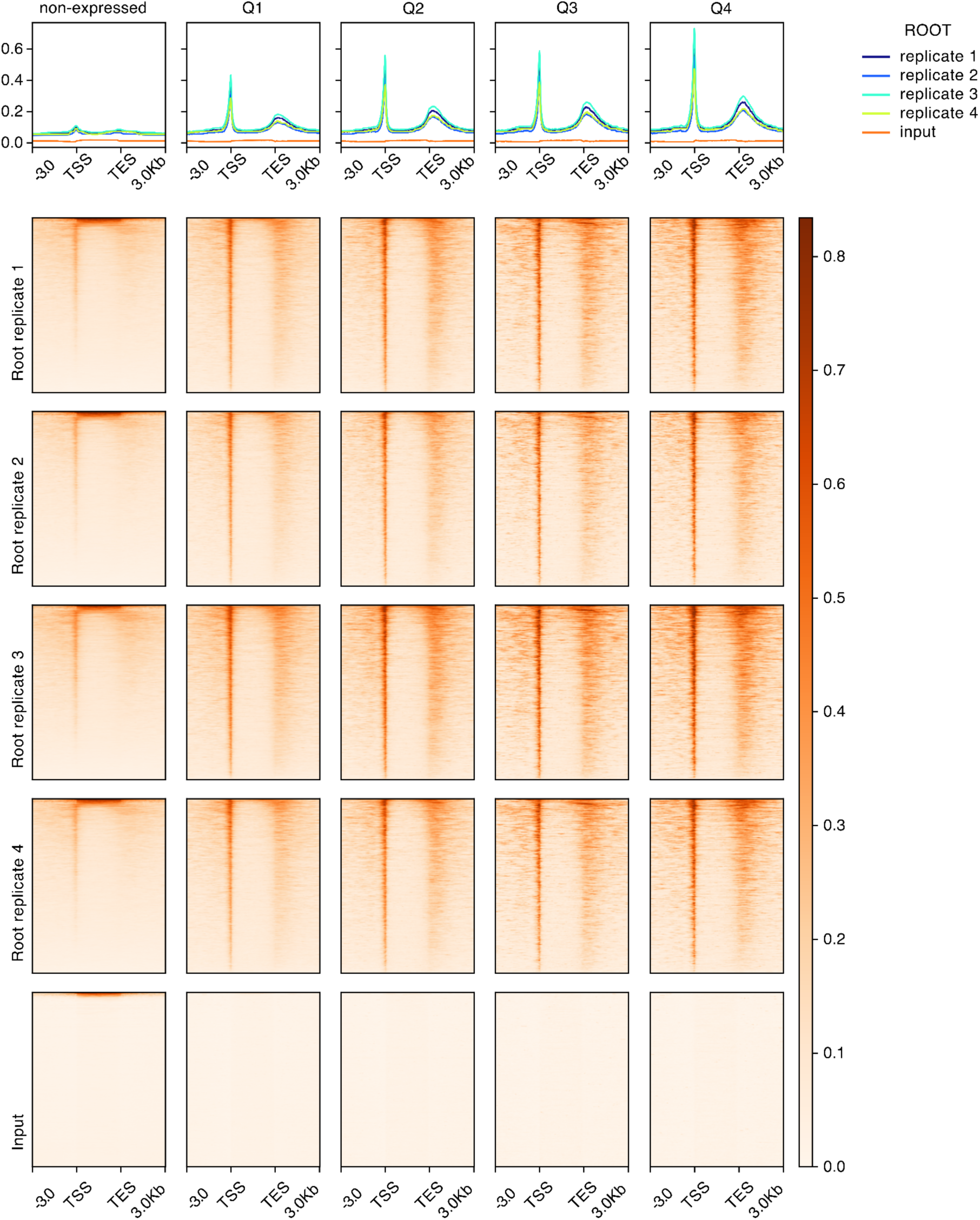
ATAC-seq metaplots of root samples. Metaplots and corresponding heatmaps of the genome-wide ATAC-seq signal (normalised by reads per million) of each leaf and input sample. Data was computed for non-expressed genes and for expressed genes divided into four quantiles of expression level (Q1=lowest to Q4=highest) (based on normalised gene expression in 2-week-old roots). The plot represents 3 kb upstream (-3.0), transcription starting site (TSS), transcription end site (TES), and 3 kb downstream of each gene (3.0 kb). The gene body region was scaled to 3,000 bp. The profile plot represents the median of all the values for each group of genes.

**Supplementary Figure 7.**
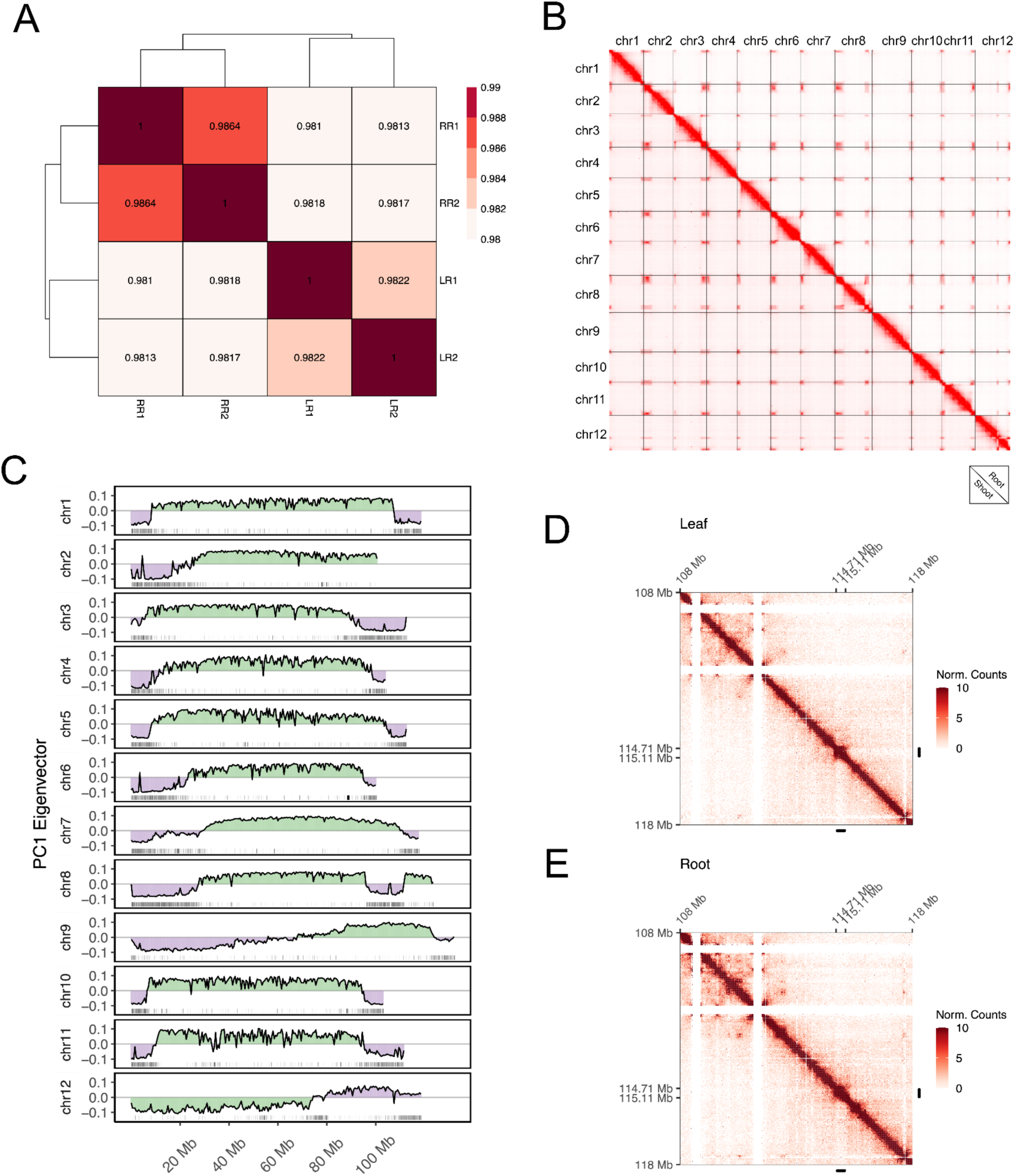
Chromosomal conformation capture by Hi-C in *P. grisea*. **A)** Hierarchical clustering calculated from the Pearson’s correlation of normalised Hi-C contact matrixes at 20-kb resolution from each sample: leaf replicates 1 and 2 (LR1, LR2); root replicates 1 and 2 (RR1, RR2). **B)** Genome-wide contact matrix of *P. grisea* at 1 Mb resolution, with shoot interactions displayed on the lower diagonal and root interactions on the upper diagonal. The colour intensity indicates frequency of interaction between two 1-Mb loci. **C)** A/B compartment annotation of *P. grisea* chromosomes. Negative (A compartment) and positive (B compartment) values represent the eigenvalues of the first principal component (PC) derived from the correlation coefficient matrix of the Observed/Expected Hi-C map for each chromosome at a 500-kb resolution. Gene density is represented at the bottom of each graph (black line). **D)** and **E)** Normalised Hi-C contact matrixes at 20-kb resolution covering the end of chromosome 1 (10 Mb) of leaf (**D)** and root (**E)** samples. Vertical and horizontal black lines indicate the withanolides BGC and denote reduced interaction with neighbouring chromatin.

**Supplementary Figure 8.**
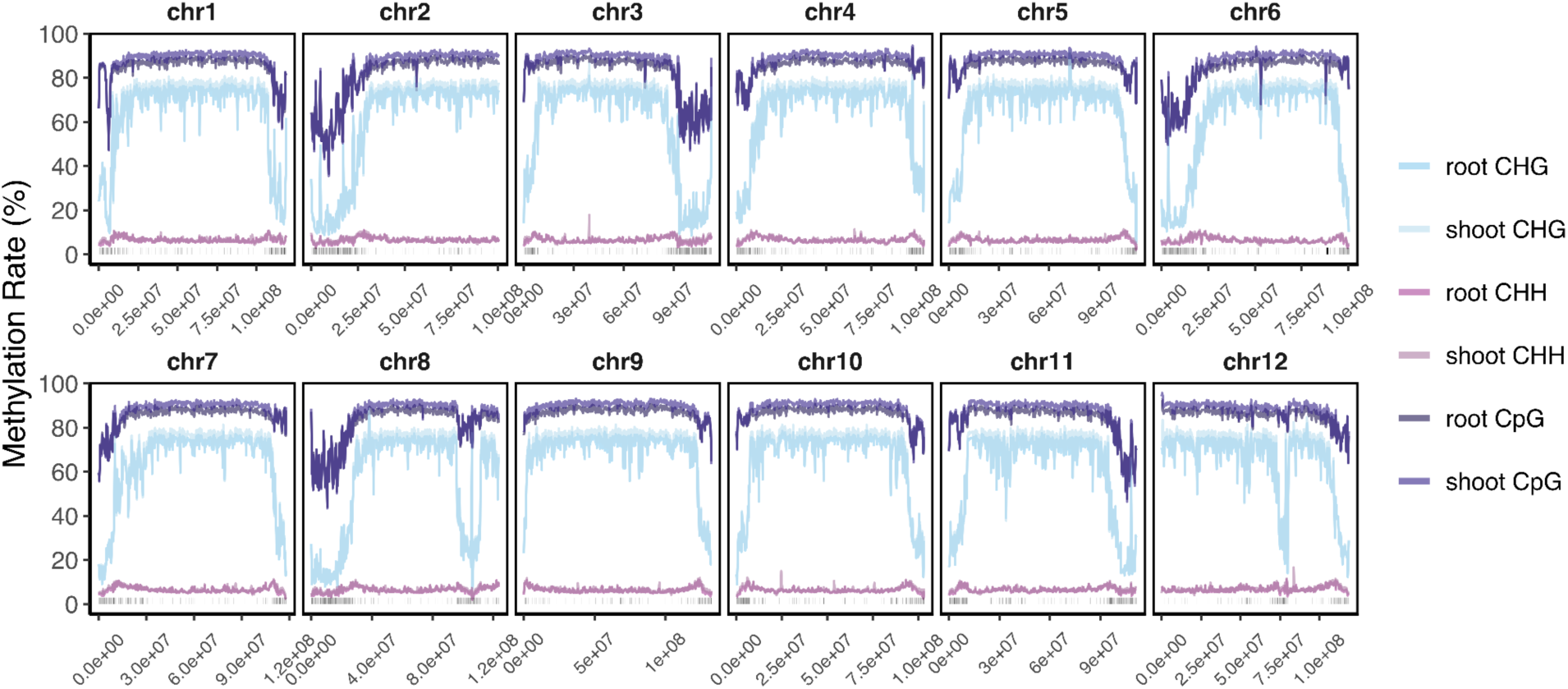
DNA methylation in *P. grisea*. Genome-wide methylation distribution in the *P. grisea* genome, calculated as the average per-site methylation rate in 300 kb windows and averaged for the three replicates in each tissue. Dark lines above the x axis represent gene models.

**Supplementary Figure 9.**
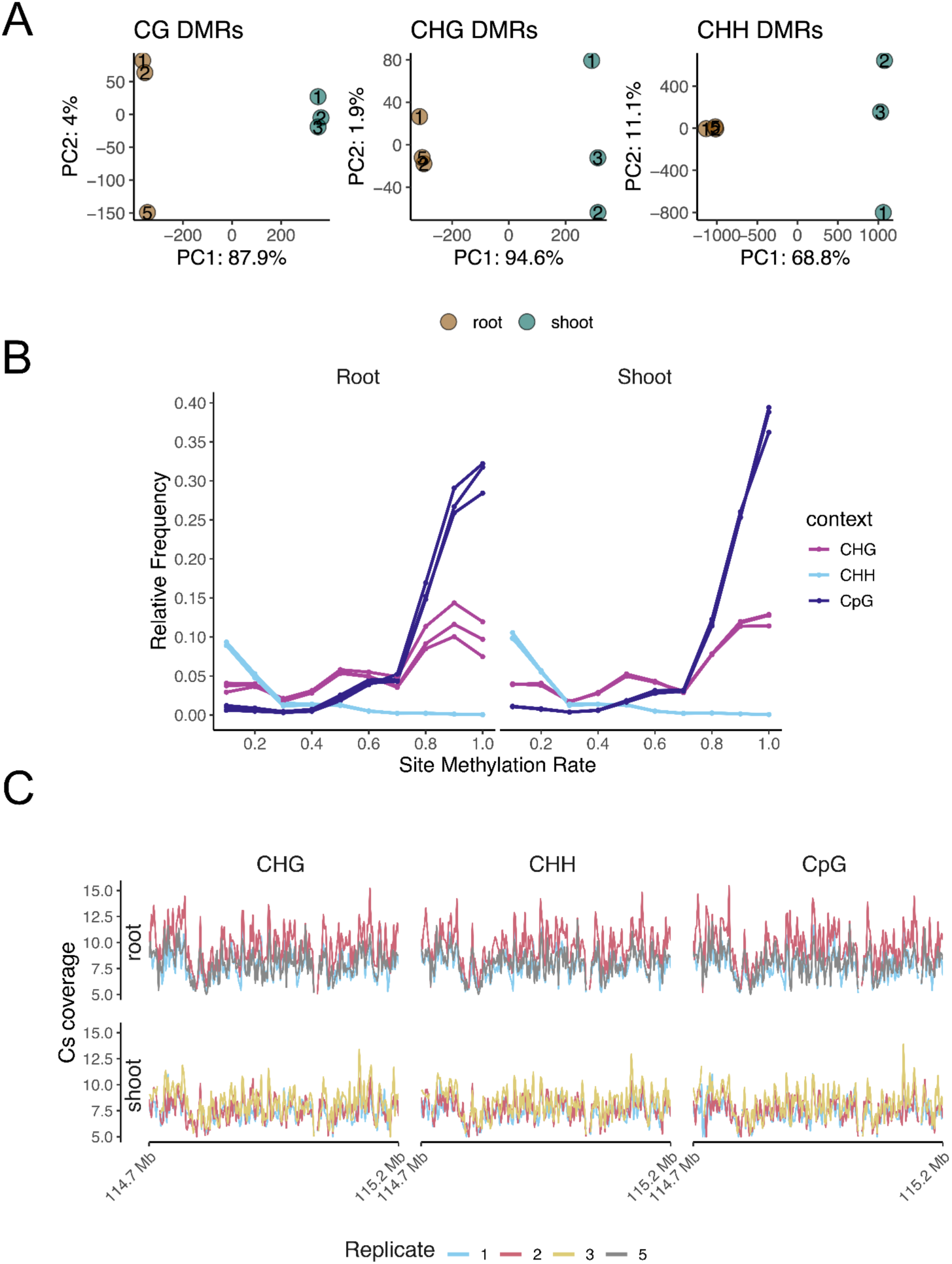
WGBS quality control. **A)** Principal components analysis of the mean methylation rate in differentially methylated regions (DMRs) detected with MethylSore (Hüther et al. 2022). Number of DMRs: CG-DMRs = 322, CHG-DMRs = 183, CHH-DMRs = 9,522. **B)** Relative frequency of the site methylation rates in *P. grisea*. Each line per context represents one replicate. **C)** Mean cytosine coverage per context in 1 kb non-overlapping windows in the withanolide BGC locus.

**Supplementary Figure 10.**
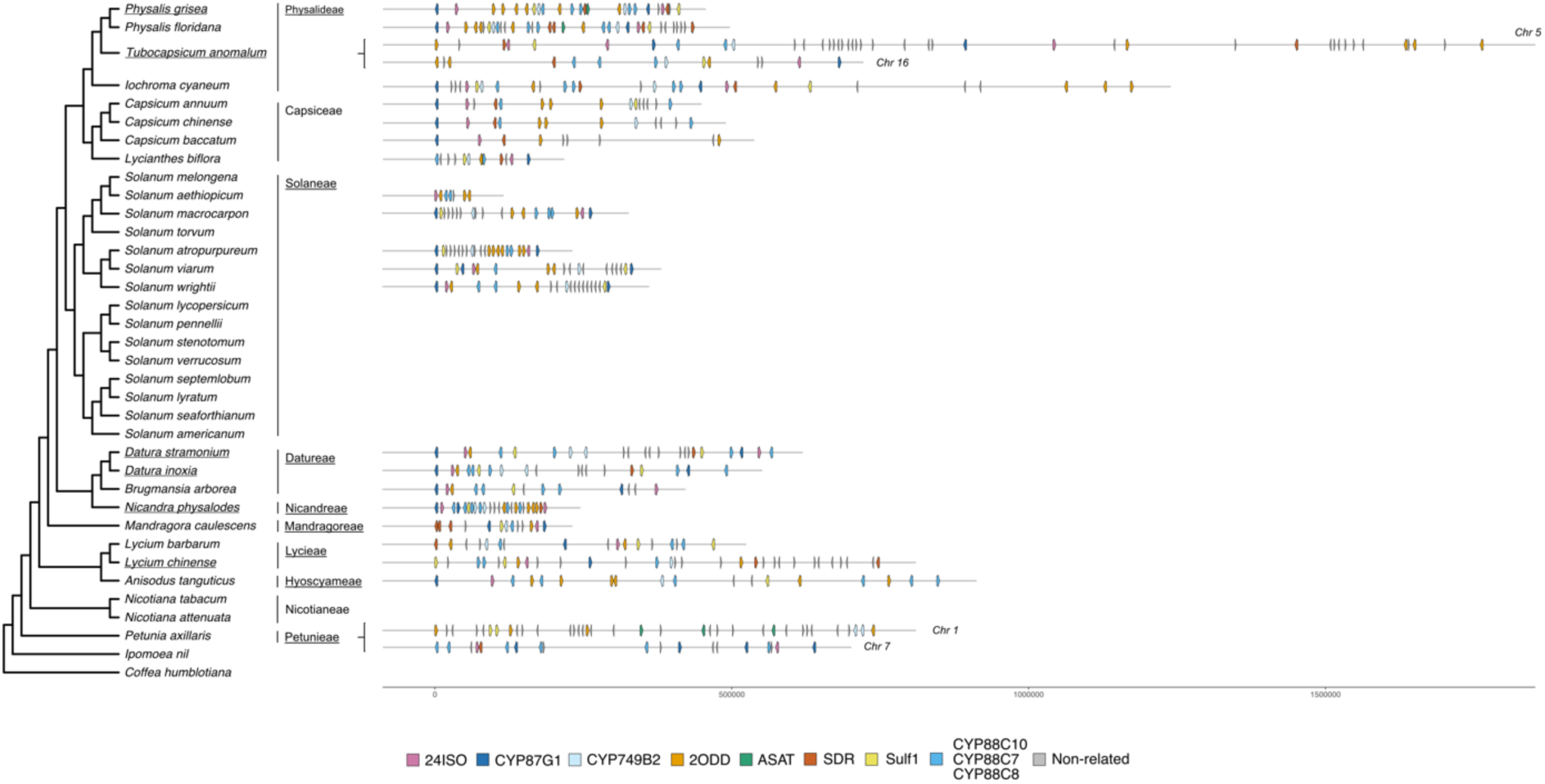
Phylogenomic comparison of the withanolide BGC across *Solanaceae*, drawn to scale. The phylogenetic tree and cluster architectures are identical to the ones shown in Figure 5. Distances between genes are drawn to scale in this version.

**Supplementary Figure 11.**
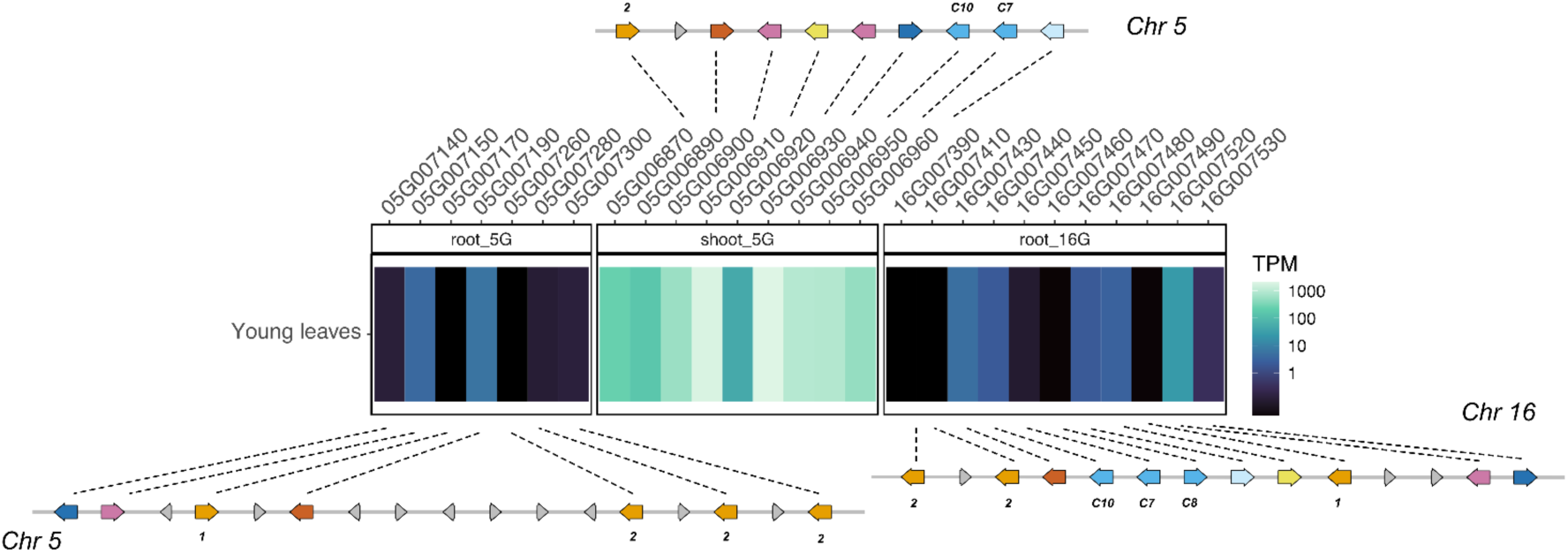
Expression of an orthologous BGC in *T. anomalum*. Expression is represented as transcript per million (TPM). Dashed lines connect each gene model with its respective expression data track.

**Supplementary Figure 12.**
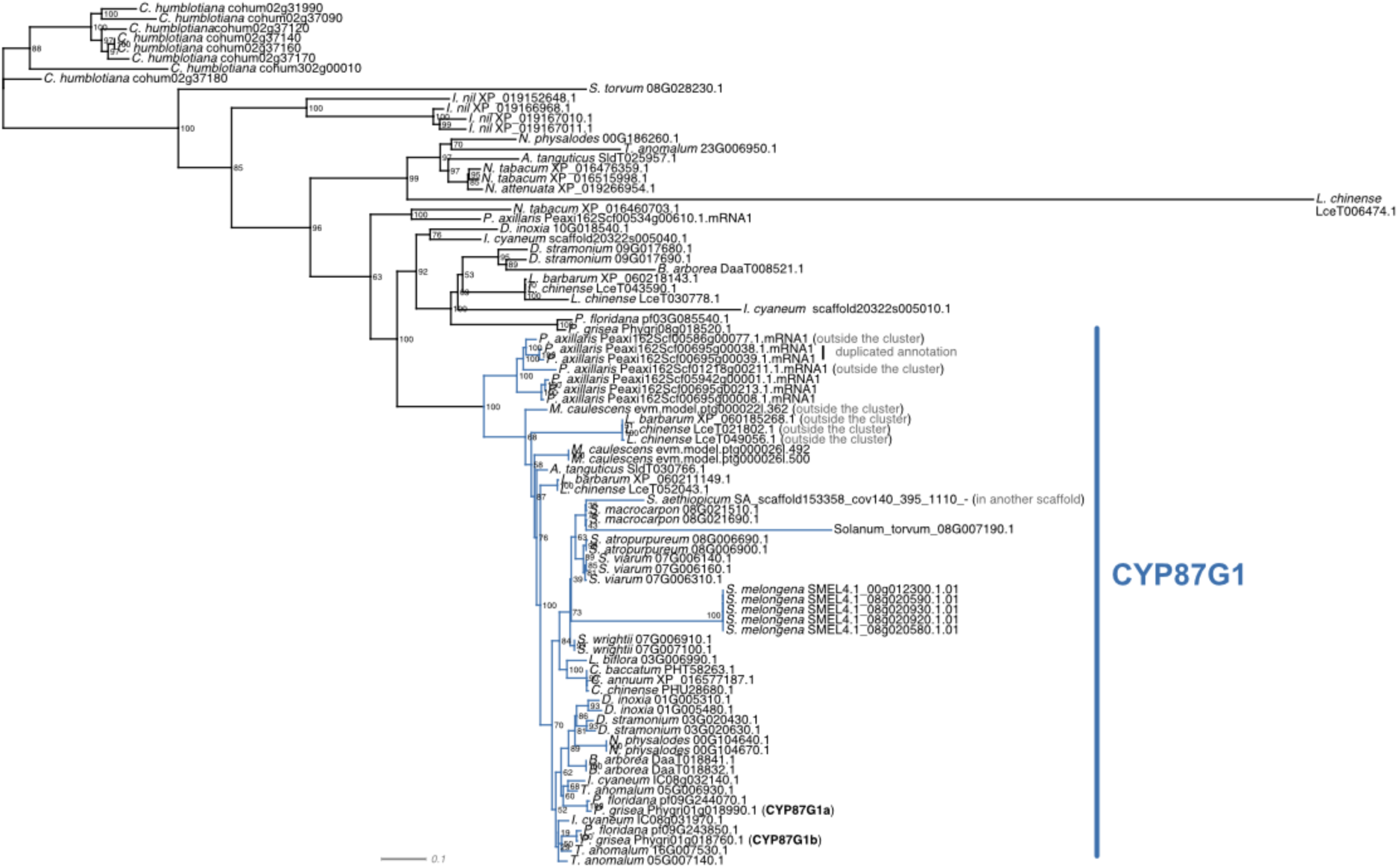
Phylogenetic analysis based on CYP87G1 amino acid sequences. Maximum likelihood tree with the best-fit model selected by ModelFinder for the orthogroup containing *P. grisea CYP87G1* paralogs (Phygri01g018760 and Phygri01g018990) in the *Solanaceae* species included in **Supplementary Table 2**. Numbers represent 1,000 replicates of ultrafast bootstrapping. Coloured edges highlight the CYP87G1 clade, which contains orthologs potentially related (but located outside the cluster) or clustered within orthologous *Solanaceae* withanolide BGCs. In the case of *P. axillaris*, duplicated annotations are indicated.

**Supplementary Figure 13.**
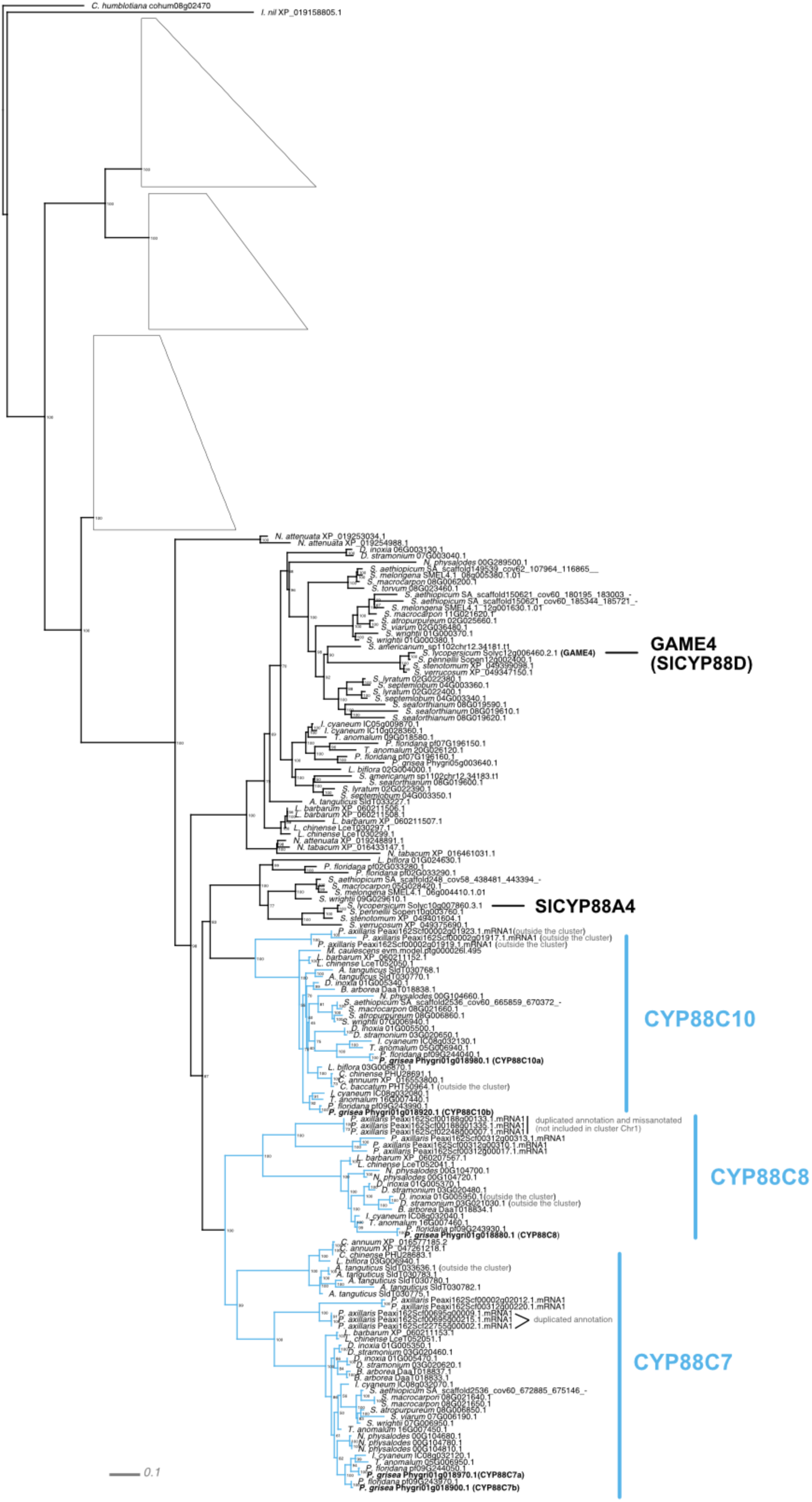
Phylogenetic analysis based on the CYP88C subfamily members (CYP88C7, CYP88C8 and CYP88C10) amino acid sequences. Maximum likelihood tree with the best-fit model selected by ModelFinder for the orthogroup containing *P. grisea CYP88C7, CYP88C8* and *CYP88C10* orthologs and paralogs (Phygri01g018880, Phygri01g018900, Phygri01g018920, Phygri01g018970 and Phygri01g018980) in the *Solanaceae* species included in **Supplementary Table 2**. Numbers represent 1,000 replicates of ultrafast bootstrapping. Coloured edges highlight the CYP88C7, CYP88C8 and CYP88C10 clades, which contain orthologs potentially related (but located outside the cluster) or clustered within orthologous *Solanaceae* withanolide BGCs. In the case of *P. axillaris*, duplicated annotations are indicated.

**Supplementary Figure 14.**
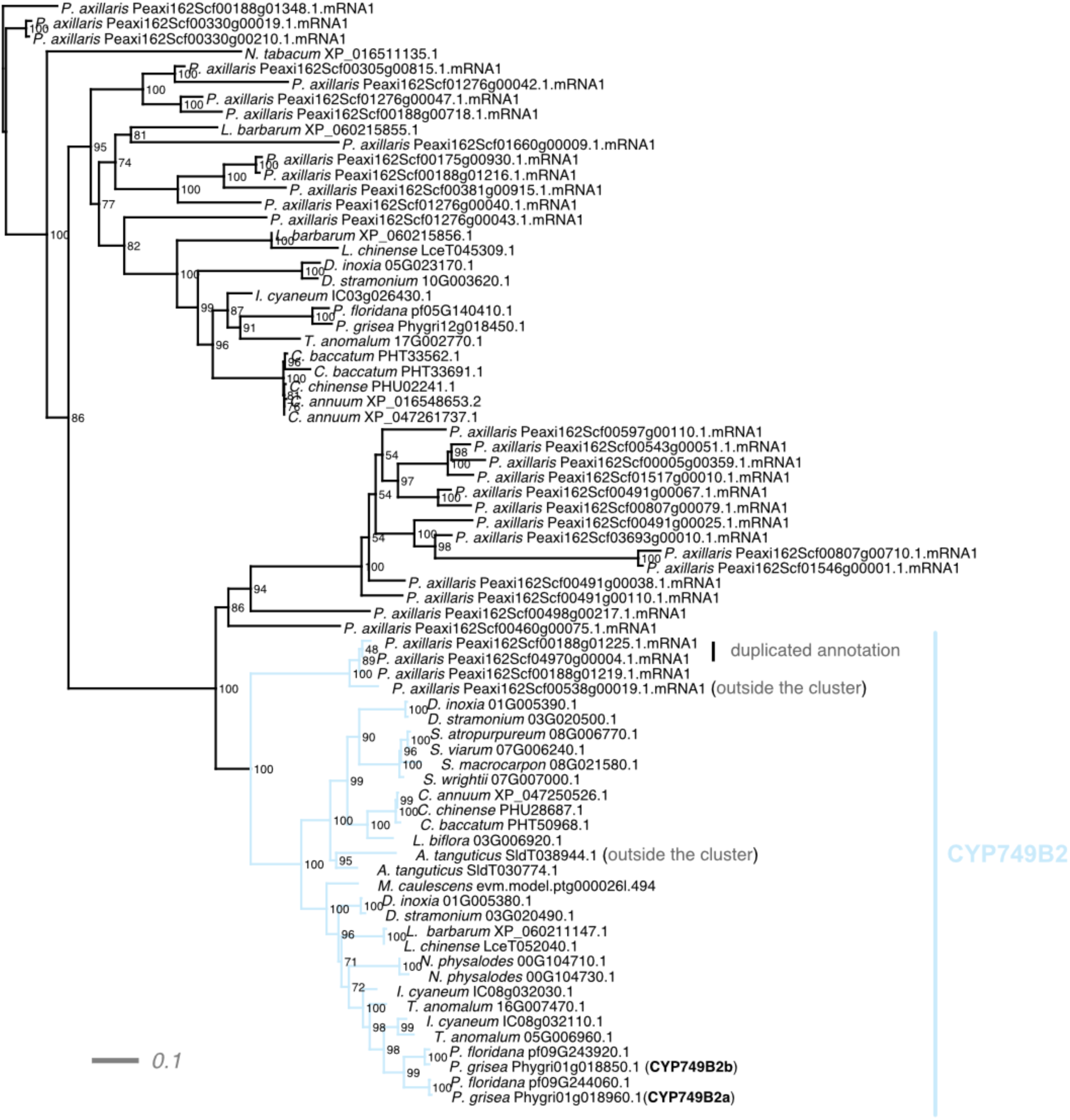
Phylogenetic analysis based on the CYP749B2 amino acid sequences. Maximum likelihood tree with the best-fit model selected by ModelFinder for the orthogroup containing *P. grisea CYP749B2* paralogs (Phygri01g018850 and Phygri01g018960) in the *Solanaceae* species included in **Supplementary Table 2**. Numbers represent 1,000 replicates of ultrafast bootstrapping. Coloured edges highlight the CYP749B2 clade, which contain orthologs potentially related (outside the cluster) or clustered in *Solanaceae* species orthologous withanolide BGCs. In the case of *P. axillaris*, duplicated annotations are indicated.

**Supplementary Figure 15.**
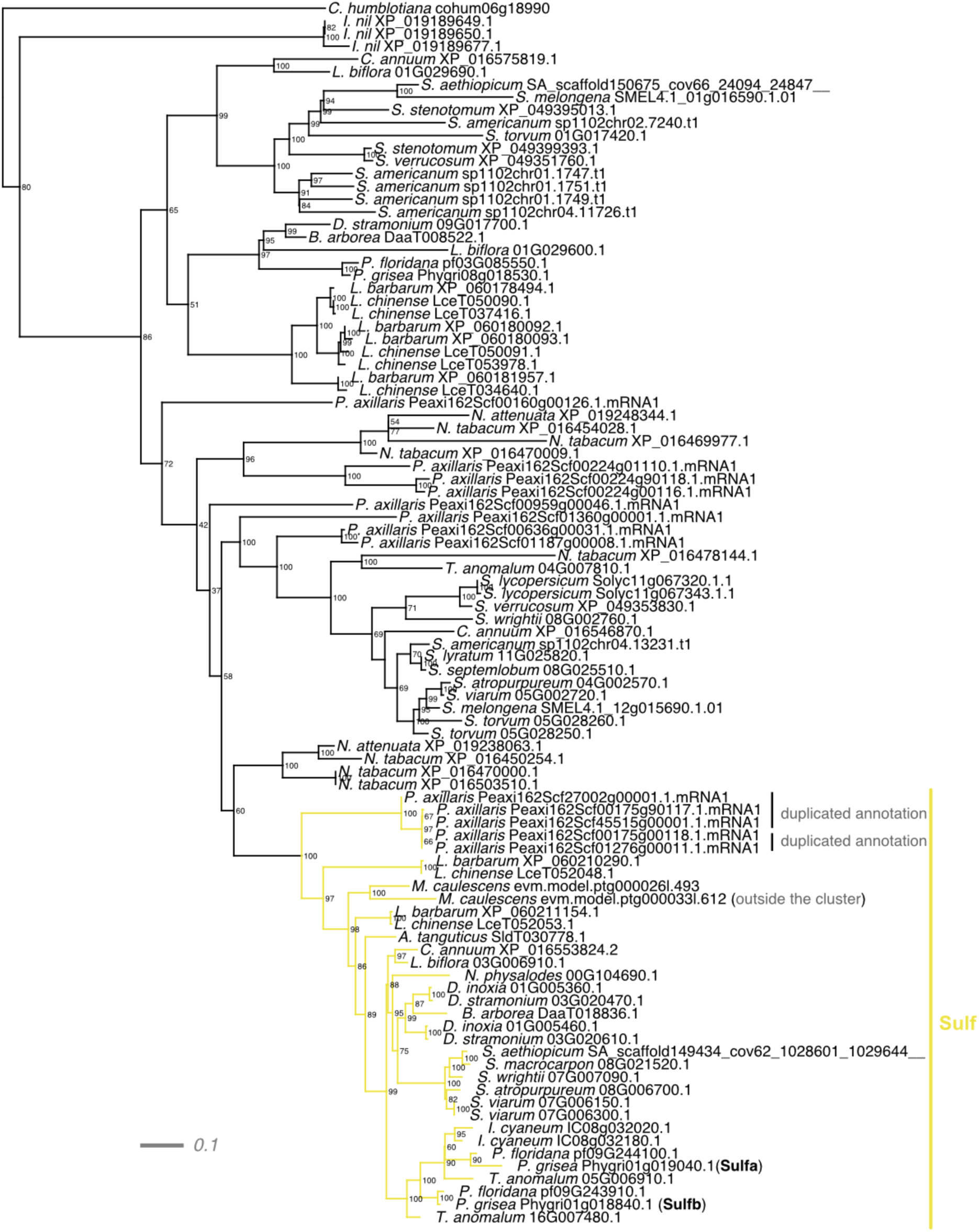
Phylogenetic analysis based on the sulfotransferases (Sulfa and Sulfb) amino acid sequences. Maximum likelihood tree with the best-fit model selected by ModelFinder for the orthogroup containing *P. grisea Sulf* paralogs (Phygri01g018840 and Phygri01g019040) in the *Solanaceae* species included in **Supplementary Table 2**. Numbers represent 1,000 replicates of ultrafast bootstrapping. Coloured edges highlight the Sulf clade, which contain orthologs potentially related (but located outside the cluster) or clustered within orthologous *Solanaceae* withanolide BGCs. In the case of *P. axillaris*, duplicated annotations are indicated.

**Supplementary Figure 16.**
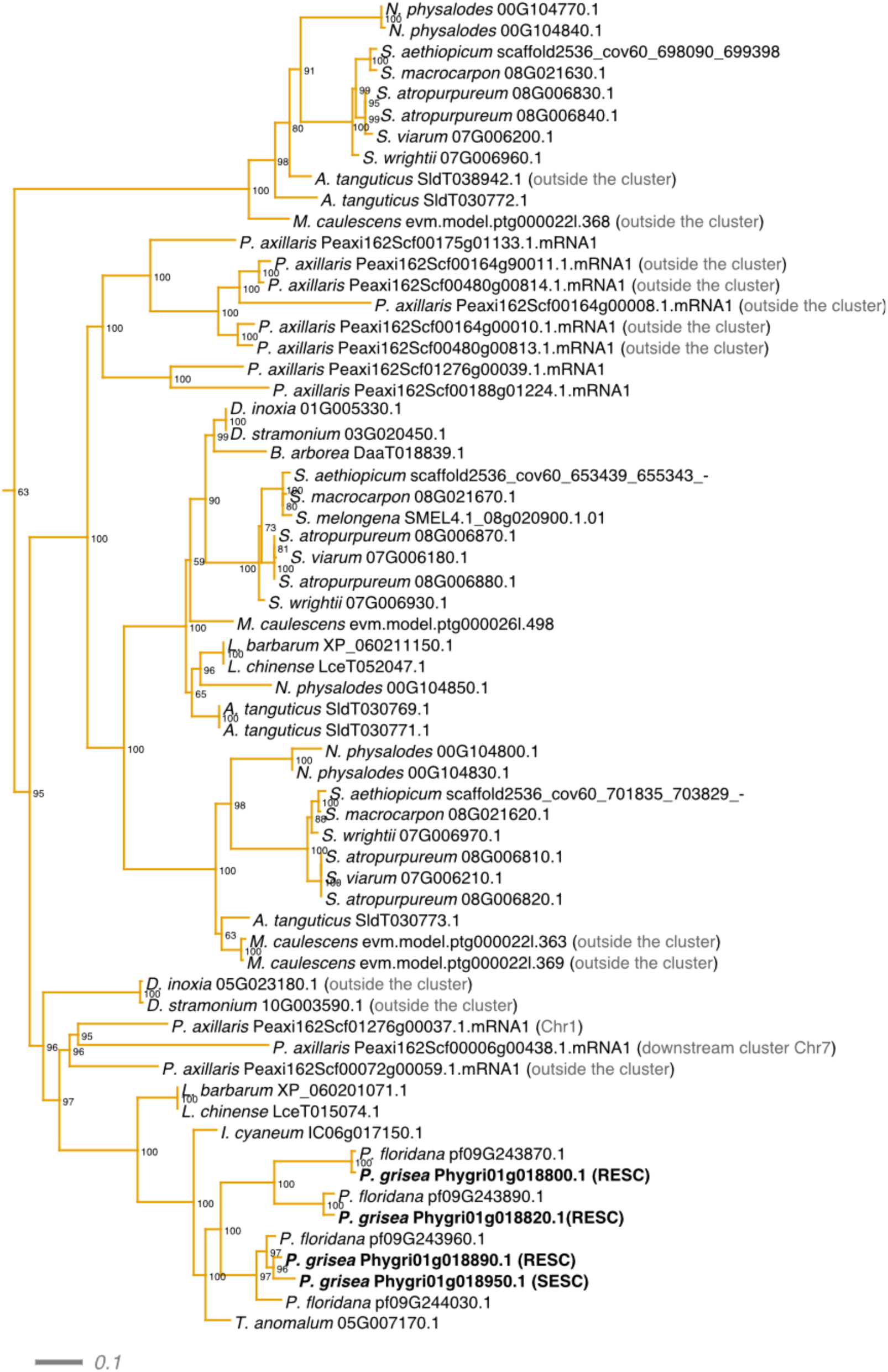
Phylogenetic analysis based on the 2ODD amino acid sequences found in the larger 2ODD orthogroup (2ODD1). Subtree from the maximum likelihood tree with the best-fit model selected by ModelFinder for the orthogroup containing *P. grisea* 2ODD1 paralogs (Phygri01g018800, Phygri01g018820, Phygri01g018890 and Phygri01g018950) in the *Solanaceae* species included in **Supplementary Table 2**. Numbers represent 1,000 replicates of ultrafast bootstrapping. Coloured edges highlight the 2ODD1 clade, which contain orthologs potentially related (but located outside the cluster) or clustered within orthologous *Solanaceae* withanolide BGCs.

**Supplementary Figure 17.**
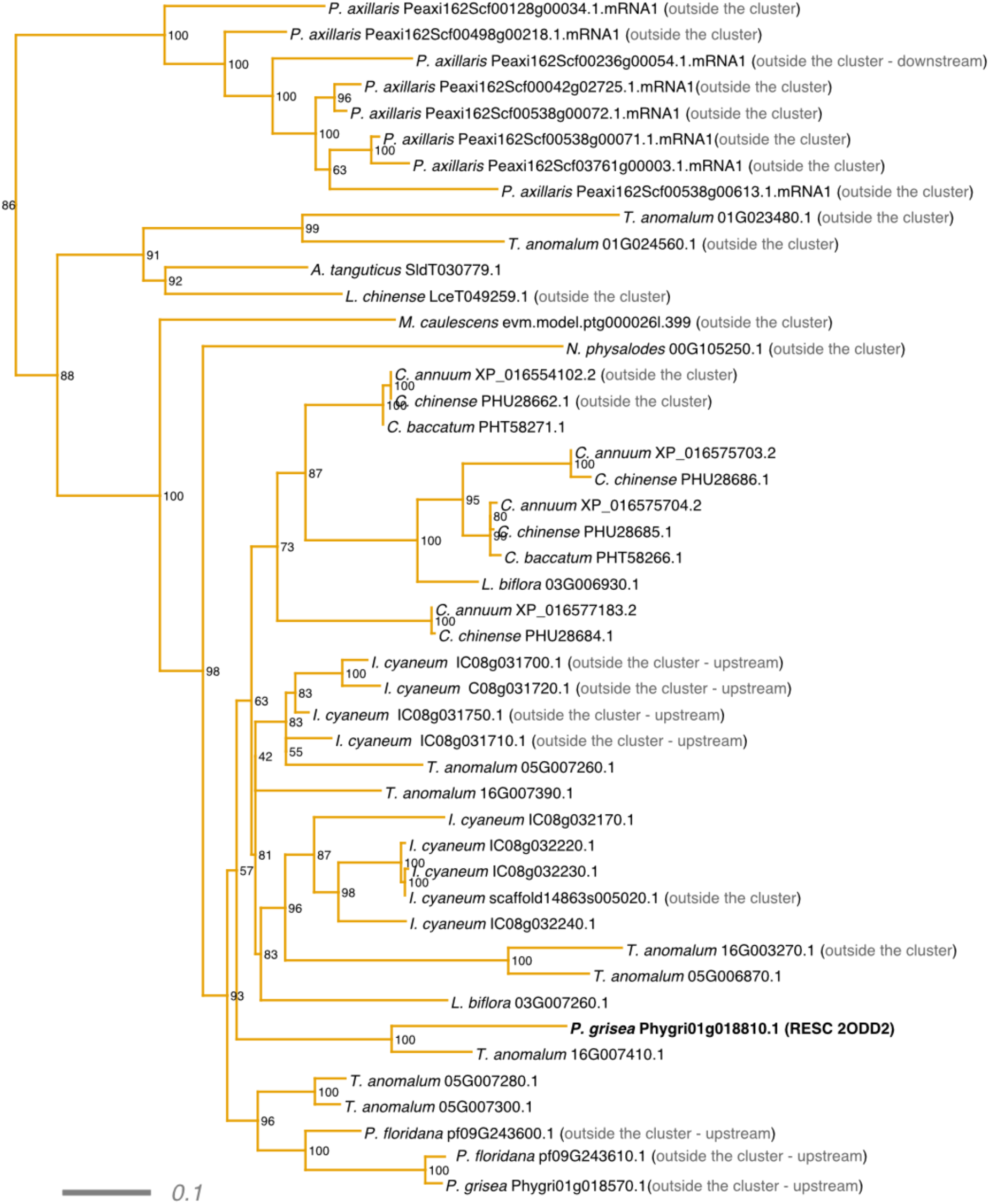
Phylogenetic analysis based on the 2ODD amino acid sequences found in the second larger 2ODD orthogroup (2ODD2). Subtree from the maximum likelihood tree with the best-fit model selected by ModelFinder for the orthogroup containing *P. grisea* 2ODD2 (Phygri01g018810) in the *Solanaceae* species included in **Supplementary Table 2**. Numbers represent 1,000 replicates of ultrafast bootstrapping. Coloured edges highlight the 2ODD2 clade, which contain orthologs potentially related (but located outside the cluster) or clustered within orthologous *Solanaceae* withanolide BGCs.

**Supplementary Figure 18.**
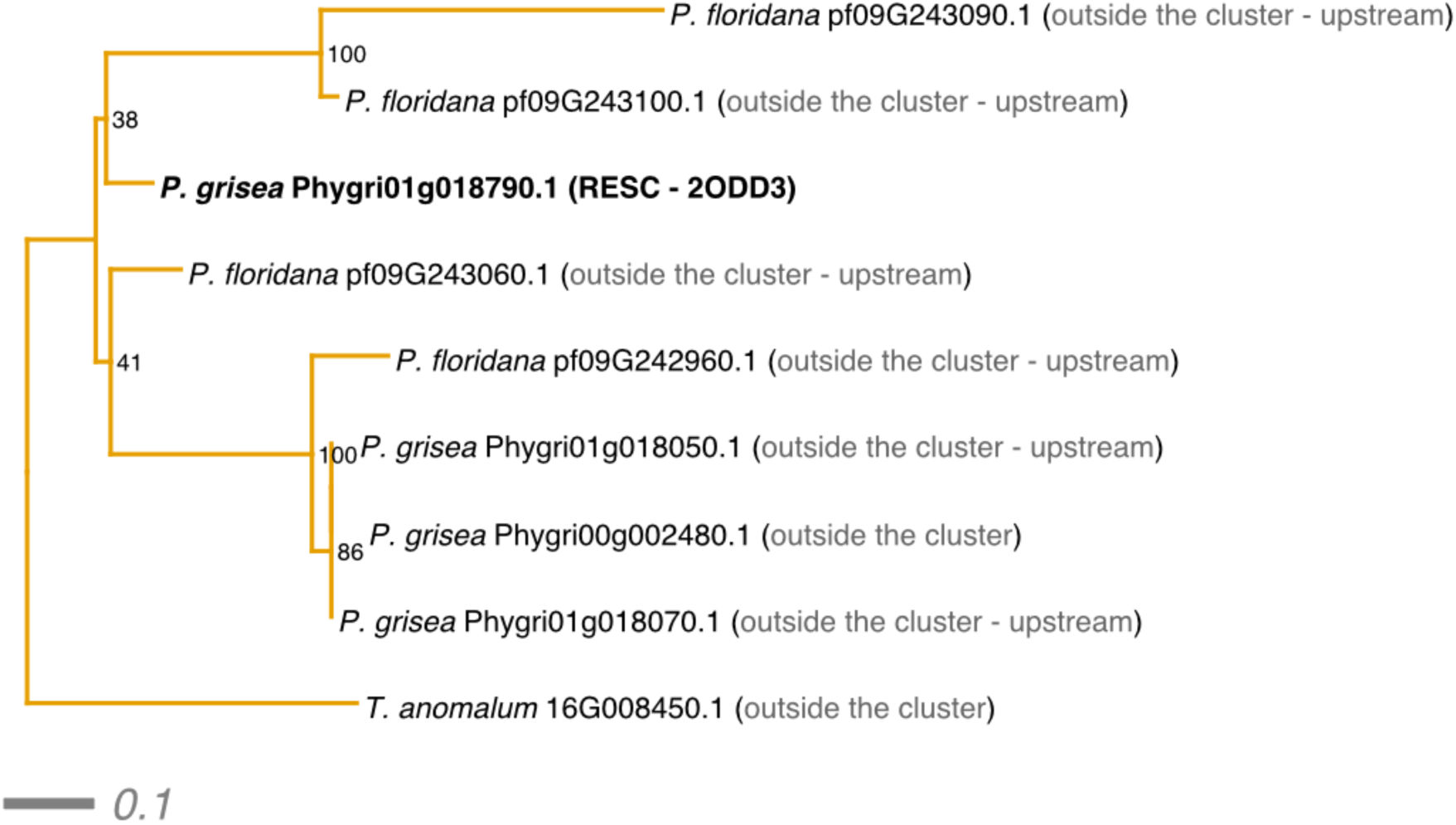
Phylogenetic analysis based on the 2ODD amino acid sequences found in the smaller 2ODD orthogroup (2ODD3). Maximum likelihood tree with the best-fit model selected by ModelFinder for the orthogroup containing *P. grisea* 2ODD3 (Phygri01g018790) in the *Solanaceae* species included in **Supplementary Table 2**. Numbers represent 1,000 replicates of ultrafast bootstrapping. Coloured edges highlight the 2ODD3 clade, which contain orthologs potentially related (but located outside the cluster) or clustered within orthologous *Solanaceae* withanolide BGCs.

**Supplementary Figure 19.**
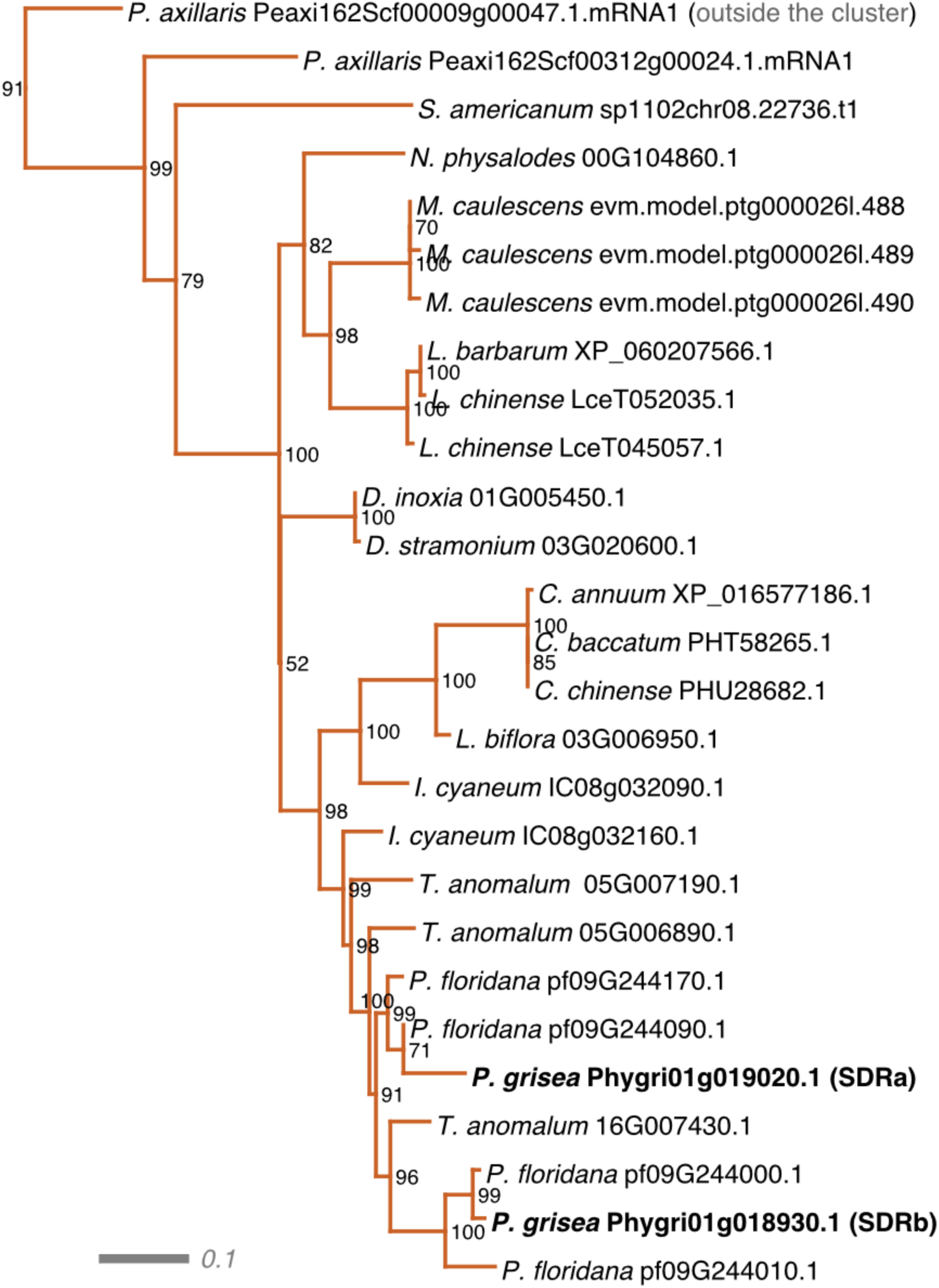
Phylogenetic analysis based on the SDRs (SDRa and SDRb) amino acid sequences. Subtree from the maximum likelihood tree with the best-fit model selected by ModelFinder for the orthogroup containing *P. grisea* SDRs (Phygri01g018930 and Phygri01g019020) in the Solanaceae species included in **Supplementary Table 2**. Numbers represent 1,000 replicates of ultrafast bootstrapping. Coloured edges highlight the SDRs clade, which contain orthologs potentially related (but located outside the cluster) or clustered within orthologous *Solanaceae* withanolide BGCs.

**Supplementary Figure 20.**
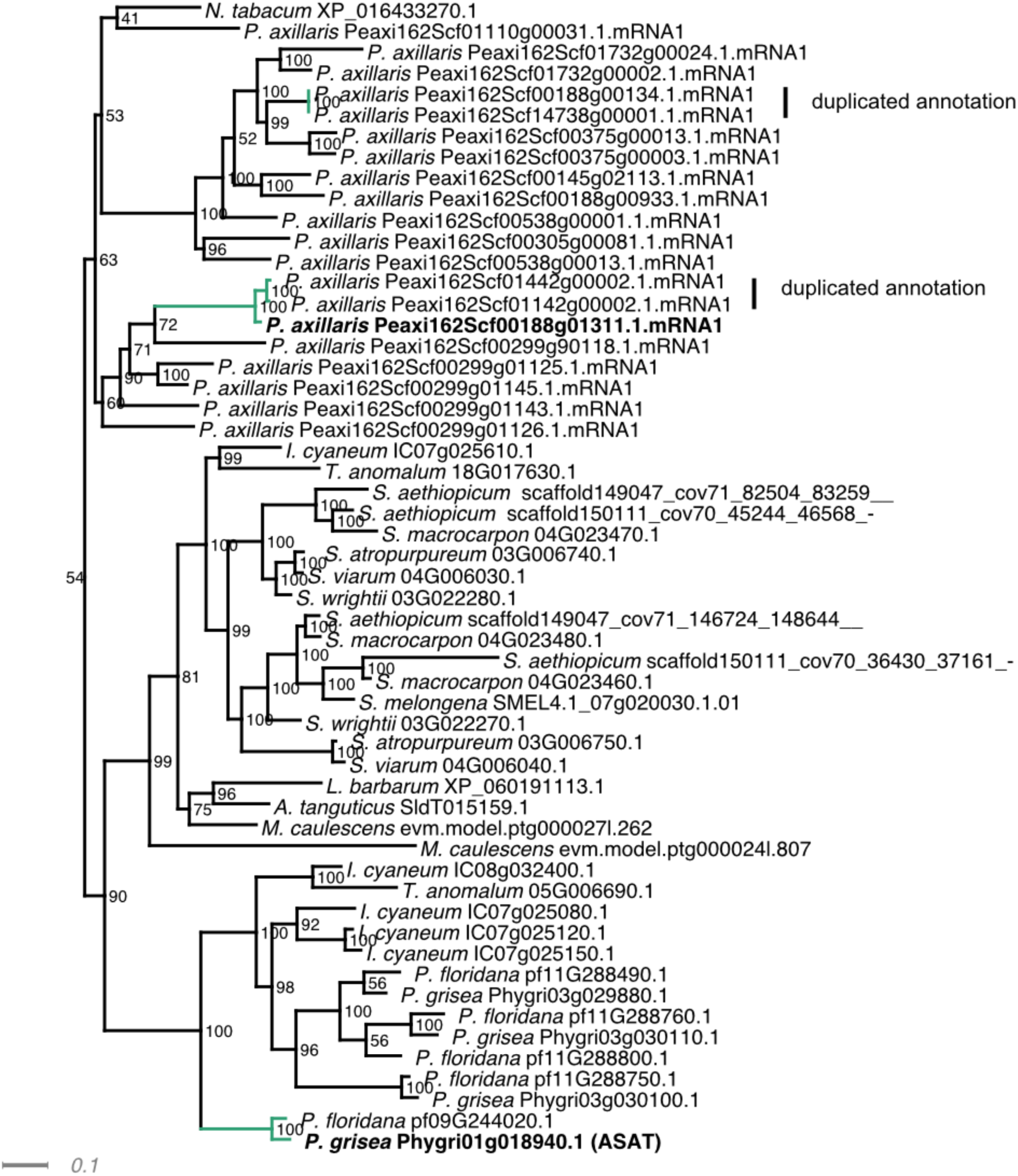
Phylogenetic analysis based on the ASAT amino acid sequences. Subtree from the maximum likelihood tree with the best-fit model selected by ModelFinder for the orthogroup containing *P. grisea* ASAT (Phygri01g018940) in the *Solanaceae* species included in **Supplementary Table 2**. Numbers represent 1,000 replicates of ultrafast bootstrapping. Coloured edges highlight the ASAT clustered in orthologous *Solanaceae* withanolide BGCs.

**Supplementary Figure 21.**
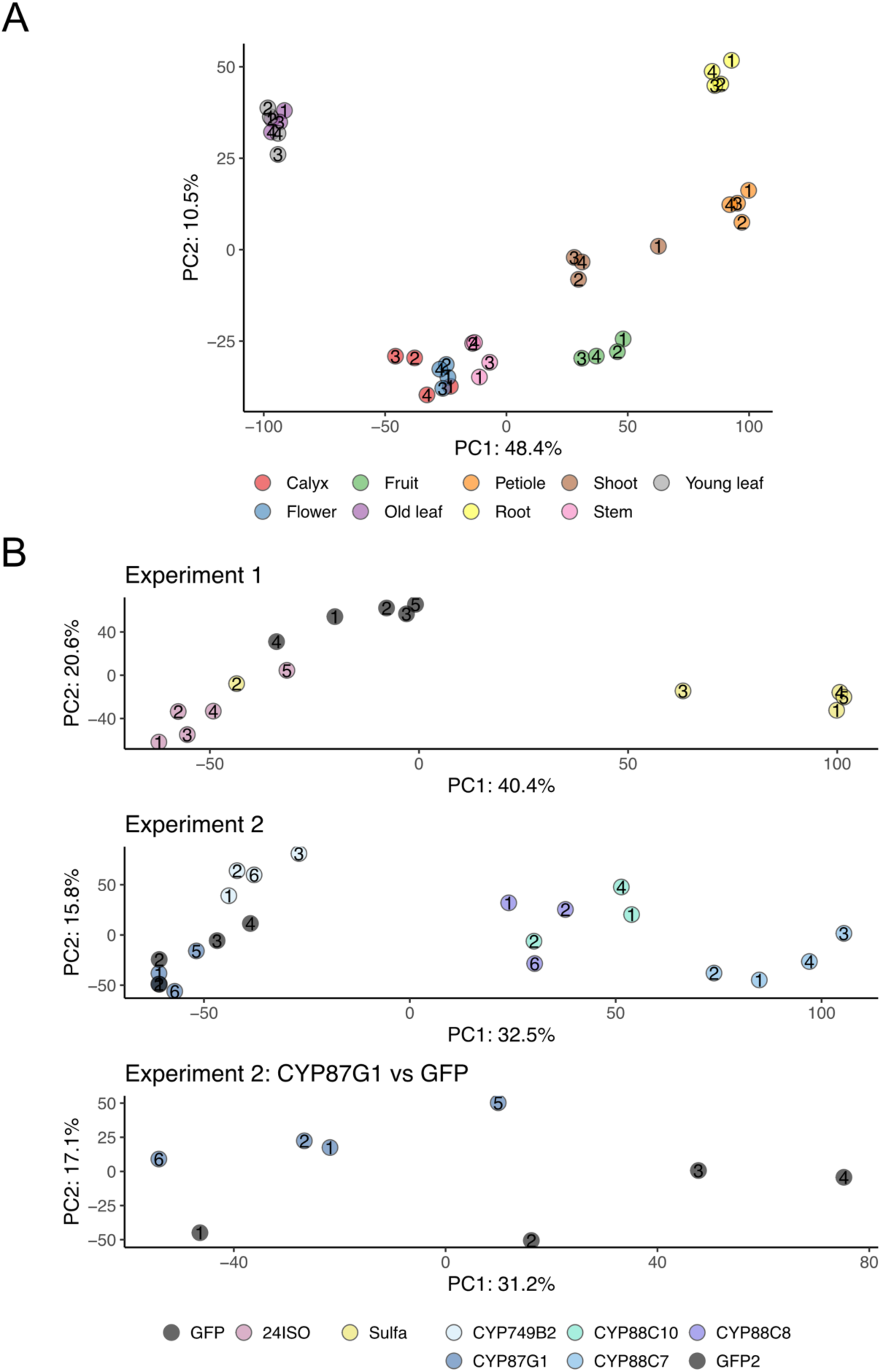
Quality control analysis of the metabolomic datasets. **A)** and **B)** Principal component analysis of metabolite profiles from the different tissues **(A)** and the VIGS-transformed samples **(B)**. For each panel, the 4th quartile of the features with the highest variance (calculated from the log2 abundance) were considered. In **B),** ‘Experiment 1’ and ‘Experiment 2’ indicate independent experiments with their respective control (GFP and GFP2 for experiments 1 and 2, respectively).

## Supplementary Tables

**Supplementary Table 1.** Sequence identities and similarities between paralogs within the withanolide biosynthetic gene cluster in *P. grisea*.

|  | CDS |  | Protein |  |
| --- | --- | --- | --- | --- |
| Pair | % identity | % similarity | % identity | % similarity |
| 24ISO | 91.9 | 91.9 | 94.9 | 97.9 |
| CYP749B2 | 94.4 | 94.4 | 91.6 | 97.7 |
| CYP87G1 | 93.5 | 93.5 | 92.8 | 97.1 |
| CYP88C7 | 94.6 | 94.6 | 95 | 96.7 |
| CYP88C8 |  |  |  |  |
| CYP88C10 | 84.4 | 84.4 | 80 | 89 |
| Sulf | 92.2 | 92.2 | 89.9 | 93.6 |

**Supplementary Table 2.**
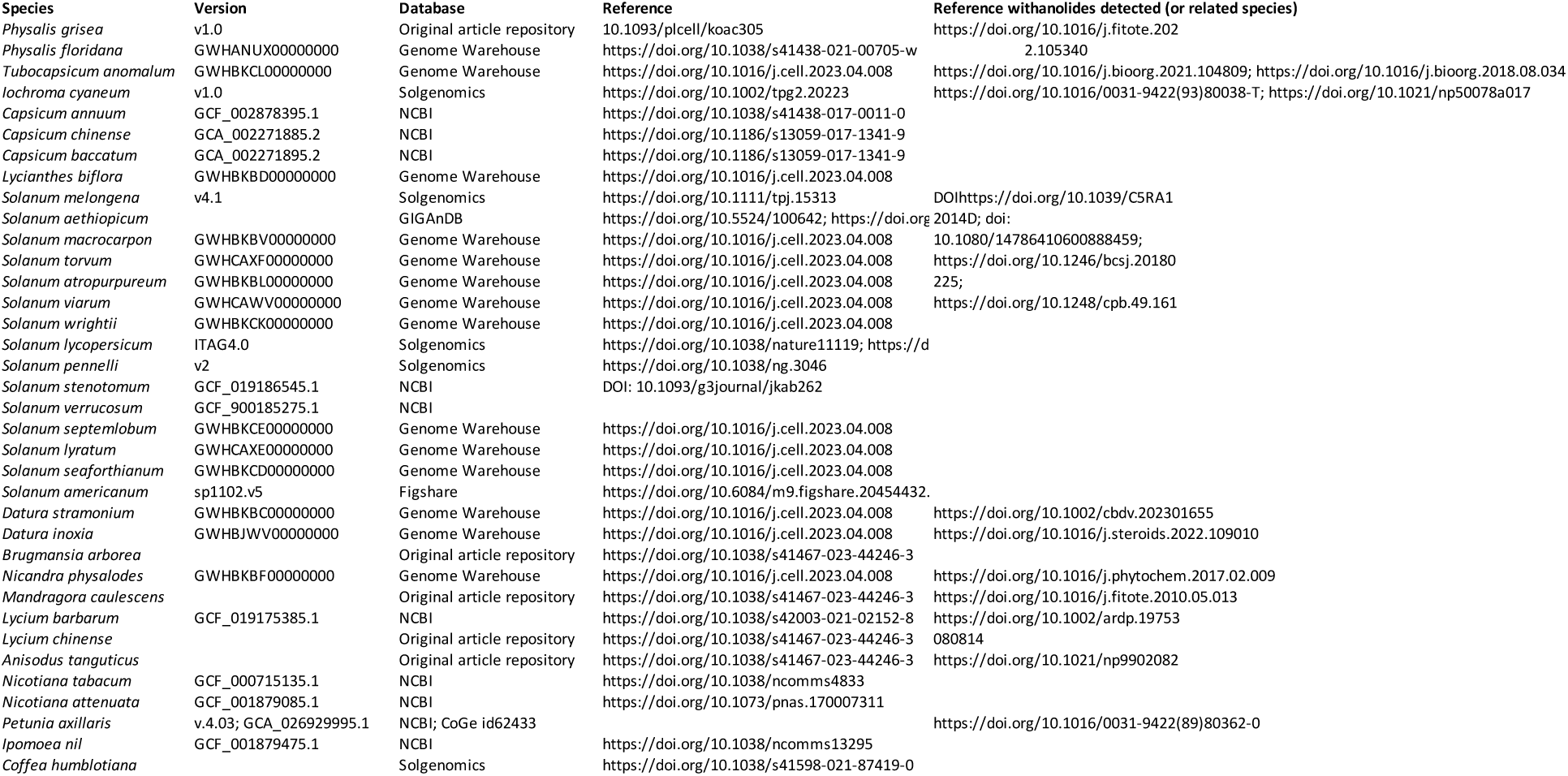
Genome assemblies used for genome mining of withanolide biosynthesis genes.

**Supplementary Table 3.**
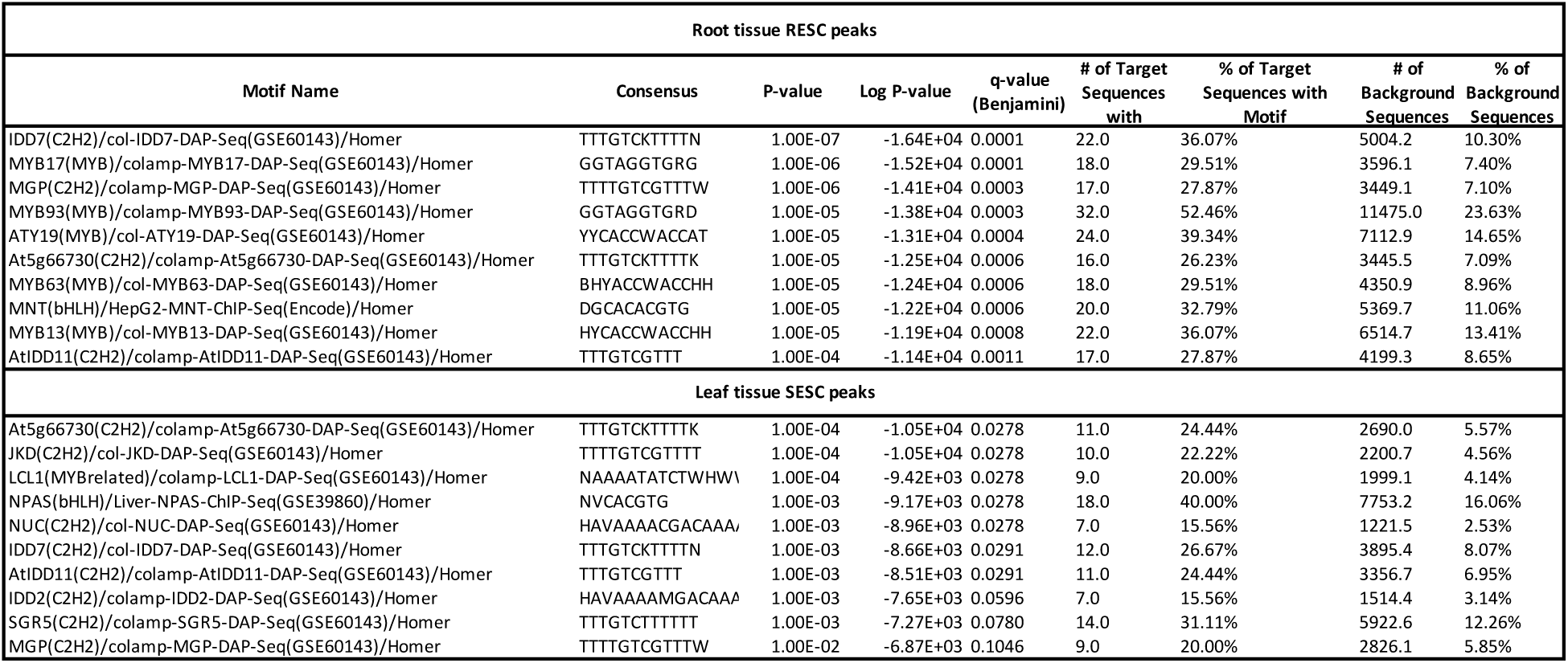
Top 10 Homer Known Motif Enrichment Results for RESC and SESC peaks in root and leaf, respectively.

**Supplementary Table 4.**
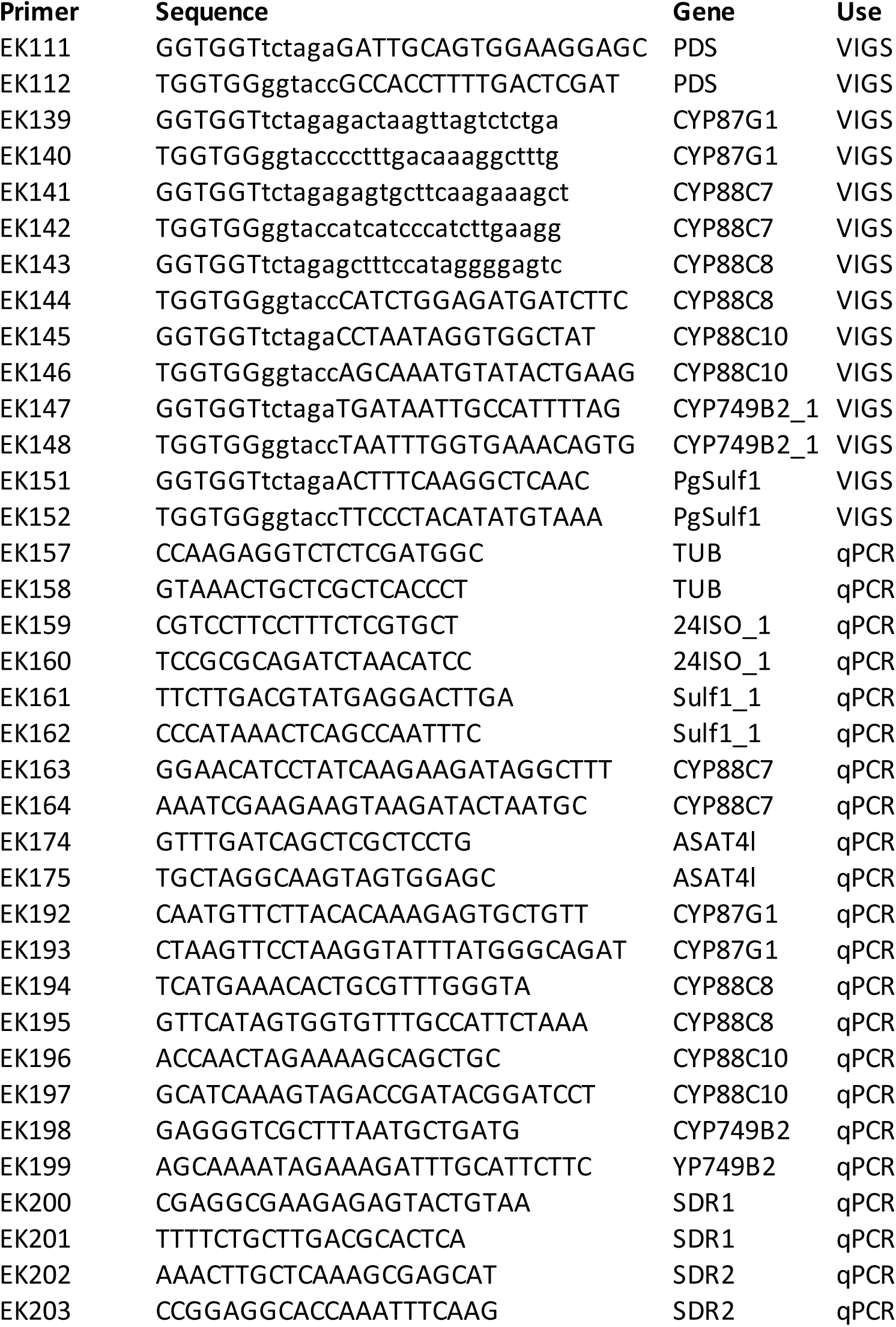
Primers used for VIGS and qPCR.

